# Lipid mobilization establishes metabolic tolerance and prevents autonomic collapse in infection

**DOI:** 10.64898/2026.04.16.717052

**Authors:** Ankita Sarkar, Shangkui Xie, Syed Muhammad Musa Ali Rizvi, Tadiwanashe Gwatiringa, Sophie Heston, Samuel Piaker, Narges Alipanah-Lechner, Jianyi Yin, Laurent Gautron, Soumya Kamath, Neethu Alex, Ashutosh Shukla, Lin Jia, Rani Shiao, Lauren E. Kemp, David G. Thomas, Alexander M. Tatara, Catherine Chen, Mujeeb Basit, Xiaofei Kong, Vanessa Nomellini, Anoj Ilanges, Samuel Heaselgrave, Joel K. Elmquist, Heather W. Stout-Delgado, Edward J. Schenck, Angela J. Rogers, Carolyn S. Calfee, Michael A. Matthay, Shunxing Rong, Jay D. Horton, Kartik N. Rajagopalan, Suraj J. Patel

## Abstract

Survival during infection depends on both pathogen clearance and the ability to tolerate infection-induced physiological changes. Metabolic adaptations are a central component of this tolerance, but the mechanisms underlying these responses remain incompletely defined. Here, we identify white adipose tissue (WAT) lipolysis as a central regulator of metabolic tolerance to infection. In patients with sepsis, higher circulating non-esterified fatty acid (NEFA) levels were associated with reduced mortality. In mouse models of polymicrobial sepsis, infection induced robust adipose lipolysis and increased circulating NEFAs. Genetic ablation of adipose triglyceride lipase (ATGL) in adipose tissue impaired lipolysis, leading to hypothermia, bradycardia, and increased mortality without altering immune cell populations or pathogen burden, consistent with a defect in tolerance rather than resistance. Mechanistically, lipolysis-derived NEFAs, but not glycerol, were required for protection, as restoring circulating NEFAs rescued autonomic stability and survival in adipose tissue ATGL-deficient mice. Infection-induced lipolysis was redundantly regulated and did not depend on any single upstream signaling pathway. Both pharmacologic activation of lipolysis using a β3-adrenergic agonist and exogenous fatty acid supplementation increased circulating NEFAs, improved survival, and promoted tolerance in mice. Consistent with these findings, analysis of real-world electronic health record data demonstrated that septic patients receiving FDA-approved β3-adrenergic agonists had reduced mortality or hospice discharge in a propensity-matched cohort. Together, these results identify WAT lipolysis and circulating fatty acids as key mediators of tolerance to infection and support a therapeutic strategy based on repurposing clinically available β3-adrenergic agonists to improve outcomes in sepsis.

**One Sentence Summary:** White adipose tissue lipolysis promotes metabolic tolerance to infection through circulating fatty acids and is associated with improved survival in sepsis

## INTRODUCTION

Successfully overcoming infection requires both immune-mediated clearance of the pathogen and limiting collateral damage to the host, a process referred to as tolerance (1). The importance of tolerance is particularly evident in sepsis, where patients frequently develop multiorgan dysfunction and die despite timely and appropriate antimicrobial therapy (2,3). These observations demonstrate that mortality from severe infection cannot be explained solely by failure to eradicate pathogens and highlight the importance of host mechanisms that limit physiological damage during inflammatory stress. Understanding how tolerance is established during infection and how these adaptive mechanisms fail in severe disease remains a critical challenge in medicine.

Mounting an effective immune response imposes substantial metabolic demands on the host (4,5). Establishing tolerance therefore requires adapting to the energetic stresses imposed by infection (6). Activated immune cells require large amounts of energy and macromolecule substrates to support proliferation, cytokine production, and antimicrobial effector functions (7). Glucose metabolism is a major source of energy for these processes (8–10). However, infection is frequently accompanied by sickness-associated anorexia, which limits carbohydrate intake (11). Under these conditions, the host must mobilize stored energy reserves to maintain physiological function while mounting an immune response. Adipose tissue represents the largest reservoir of stored metabolic fuel and can rapidly release fatty acids through lipolysis during inflammatory stress (12–14).

Multiple infection-associated pathways have been shown to activate adipose tissue lipolysis. Adipocytes express pattern-recognition receptors (PRRs), including Toll-like receptor 4 (TLR4), which recognizes bacterial lipopolysaccharide (LPS) and can stimulate downstream ERK signaling to promote lipid mobilization (15). Inflammatory cytokines released during infection, including TNF-α, IL-1β, and IL-6, can also activate lipolytic signaling through MAPK and JAK/STAT pathways in adipocytes (16,17). In parallel, infection activates the sympathetic nervous system and increases catecholamine release, stimulating β-adrenergic receptors on white adipose tissue (WAT) to engage the canonical cAMP-PKA lipolytic pathway, characterized by phosphorylation of hormone-sensitive lipase (HSL) and activation of adipose triglyceride lipase (ATGL) (18,19). Adipose lipolysis increases circulating glycerol and non-esterified fatty acids (NEFAs). Together, these observations suggest that infection engages multiple mechanisms capable of mobilizing lipid stores. Furthermore, metabolomic profiling of patients with sepsis has identified associations between defects in fatty acid oxidation and mortality (20). However, whether adipose tissue lipolysis contributes functionally to disease tolerance during infection remains poorly understood.

To investigate the role of lipid mobilization in infection tolerance, we integrated human clinical data with genetic and pharmacologic studies in mice. In a retrospective cohort of patients with sepsis, we analyzed circulating glycerol and NEFA levels and found that higher circulating NEFA levels were associated with improved clinical outcomes, including reduced mortality. Because adipose tissue is the primary source of circulating NEFAs, we next tested whether adipose lipolysis contributes to tolerance during infection. Mice with adipocyte-specific deletion of ATGL exhibited impaired NEFA mobilization and developed severe hypothermia, bradycardia, and mortality following endotoxemia or polymicrobial sepsis, despite comparable circulating cytokine levels, immune cell activation, and systemic bacterial burden. Restoring circulating NEFAs, either by intralipid administration or pharmacologic stimulation of β3-adrenergic signaling, rescued these phenotypes, whereas glycerol supplementation did not. Consistent with these findings, analysis of nationwide electronic health record data revealed that use of β3-adrenergic agonists, including the FDA-approved drugs mirabegron and vibegron used for overactive bladder, was associated with reduced mortality among patients hospitalized with sepsis. Together, our findings establish adipose tissue lipolysis and circulating fatty acids as critical mediators of metabolic tolerance during infection.

## RESULTS

### Circulating NEFAs are associated with improved clinical outcomes in human sepsis

Circulating non-esterified fatty acids (NEFAs) were elevated in patients with infection compared to healthy individuals (**Fig. 1A**). In a cohort of critically ill patients with sepsis, analysis of plasma untargeted metabolomics data revealed that higher NEFA levels were associated with lower disease severity, as indicated by lower Acute Physiology and Chronic Health Evaluation (APACHE II) scores (**Fig. 1B**). Patient who were vasopressor-free and those who survived had significantly higher circulating NEFA levels compared to those with more severe clinical courses (**Fig. 1C, D**). Individual NEFA species, including linoleate, myristate, oleate, palmitate, and palmitoleate, were each associated with reduced risk of 60-day mortality per standard deviation increase, whereas stearate was not (**Fig. 1E**). A higher composite NEFA level was similarly associated with reduced mortality risk (aOR 0.68 per SD increase, p =0.015). Consistent with this, patients with higher composite NEFA levels exhibited improved survival, with significant separation of Kaplan-Meier curves and a reduced adjusted hazard of death (aHR 0.70, 95% CI 0.55-0.89; log-rank p = 0.010; **Fig. 1F**). In contrast to these findings, circulating glycerol levels, which are also increased during lipolysis, were not associated with sepsis severity, vasopressor requirement, or mortality (**Fig. 1G** and **Fig. S1A-D**). Together, these findings indicate that increased circulating NEFAs, but not glycerol, are associated with improved clinical outcomes in human sepsis.

**Figure 1.**
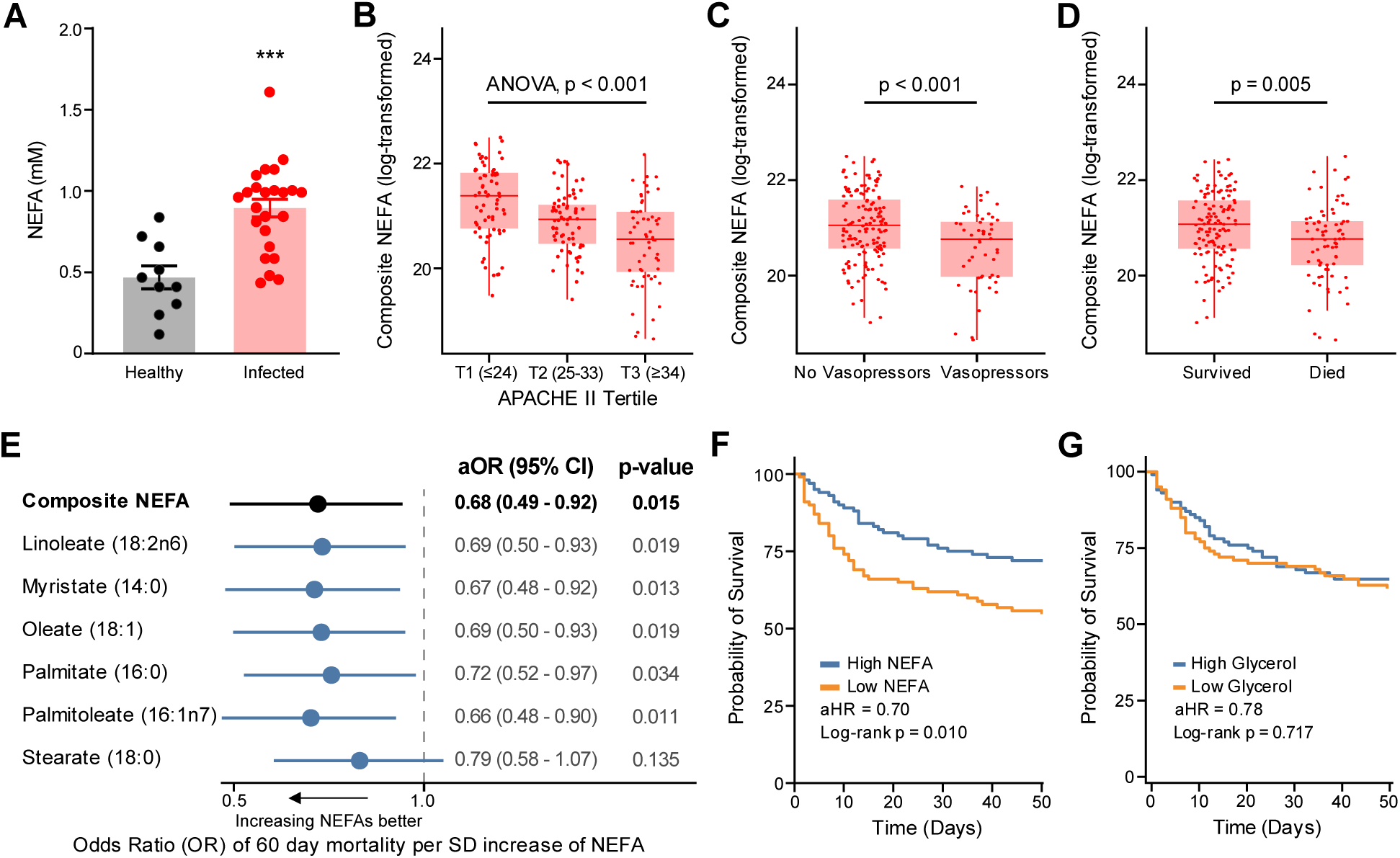
Circulating NEFAs associate with disease severity and survival in human sepsis. **(A)** Serum non-esterified fatty acid (NEFA) levels in healthy individuals and patients with infection. **(B)** Composite NEFA score stratified by Acute Physiology and Chronic Health Evaluation (APACHE II) tertiles (T1–T3). **(C)** Composite NEFA score stratified by need for vasopressors. **(D)** Composite NEFA score stratified by 60-day mortality. **(E)** Forest plot of odds ratios (ORs) for 60-day mortality from logistic regression models for each individual NEFA and the composite score, per standard deviation increase, adjusted for age, sex, and diabetes. Kaplan-Meier survival curves stratified by **(F)** composite NEFA score and **(G)** glycerol score (High vs. Low). The adjusted hazard ratio (HR) was derived from a Cox proportional hazards model adjusted for age, sex, and diabetes; the log-rank p-value for the unadjusted comparison between groups is shown. Composite NEFA score and glycerol score calculated as discussed in Materials and Methods. In panels B–D, horizontal bars indicate group means with error bars indicating standard error of the mean (SEM); significance was assessed by Wilcoxon rank-sum test (two-group comparisons) or pairwise comparison between T1 and T3; *p < 0.05, **p < 0.01, ***p < 0.001, ****p < 0.0001.

### Infection induces adipose lipolysis and a conserved metabolic stress response

To evaluate the role of lipolysis and lipid availability in infection, we measured circulating NEFA levels in models of polymicrobial sepsis and sterile inflammation. Consistent with our findings in patients with sepsis, cecal ligation and puncture (CLP) in mice increased serum NEFAs and glycerol concentrations (**Fig. 2A, B**). Serum ketone levels were also increased, indicating increased hepatic ketogenesis from mobilized fatty acids (**Fig. 2C**). Because circulating NEFAs are derived predominantly from WAT, we next examined WAT lipolytic signaling during infection. CLP activated multiple pathways known to regulate WAT lipolysis, including adrenergic-PKA, MAPK, and JAK/STAT signaling, as evidenced by increased phosphorylation of HSL, ERK1/2, STAT3, p38, and PKA substrates **(Fig. 2D).** Triglyceride lipolysis in adipocytes was accompanied by conserved physiological changes including hypothermia, bradycardia, hypoglycemia, and mortality, hallmarks of severe systemic infection (**Fig. 2E-H**). LPS-induced inflammation similarly activated lipolytic signaling in WAT and recapitulated the associated metabolic and physiologic responses found in the cecal ligation and (CLP) model (**Fig. S2**).

**Figure 2.**
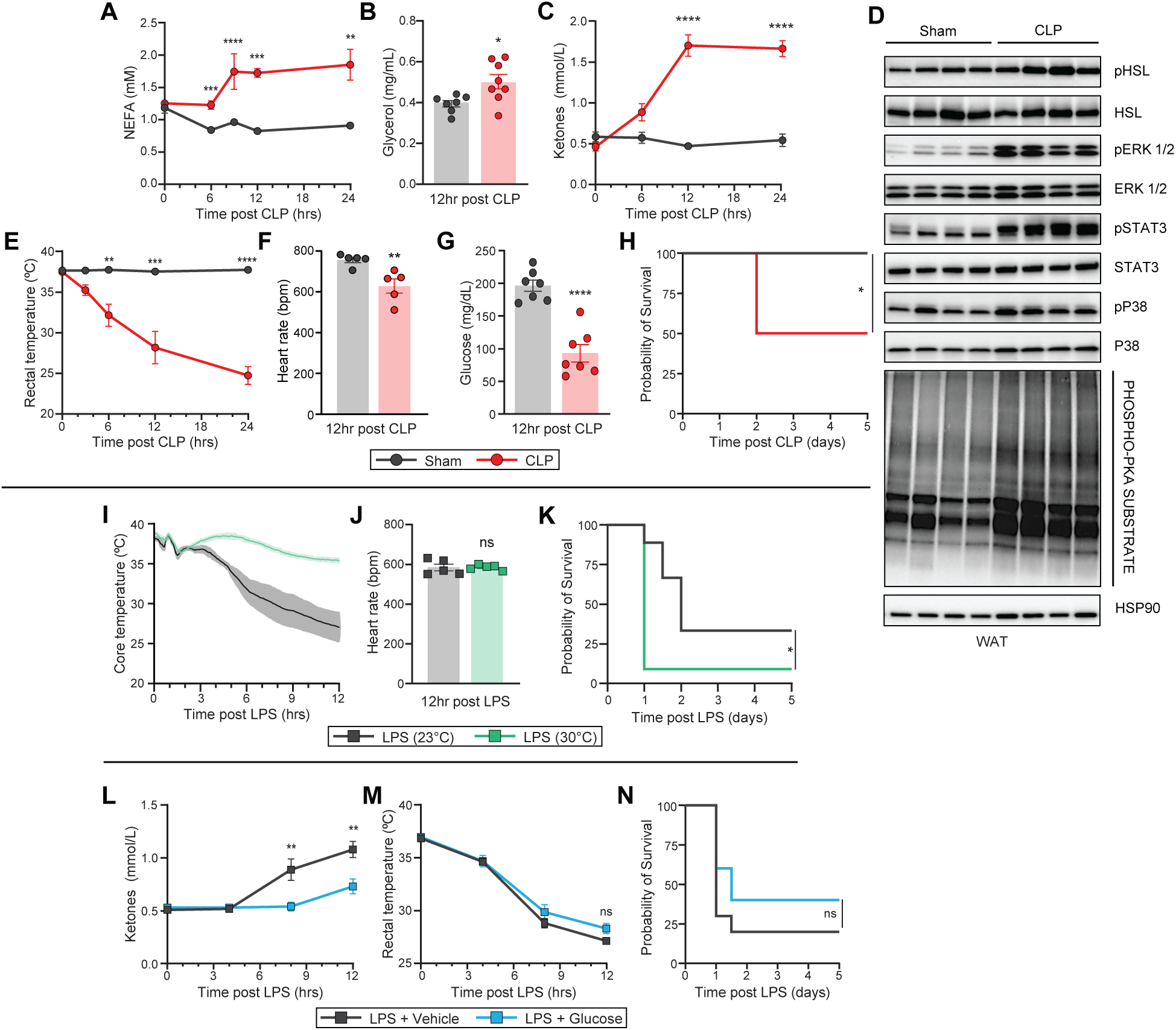
Infection induces adipose lipolysis and a conserved metabolic stress response. Circulating **(A)** non-esterified fatty acids (NEFAs) **(B)** glycerol and **(C)** ketones (n=7-8/group) after cecal ligation and puncture (CLP). **(D)** Immunoblot of pHSL, HSL, pERK, ERK, pSTAT3, STAT3, pP38, P38 and phospho-PKA substrate from white adipose tissue (WAT) after CLP (n=4/group). **(E)** Rectal temperatures (n=7-8/group) and **(F)** heart rates after CLP (n=5/group). **(G)** Circulating glucose levels, and **(H)** Kaplan–Meier survival curve after CLP (n=7-8/group). **(I)** Core temperatures (n=4-5/group), **(J)** heart rates (n=5/group) and **(K)** Kaplan–Meier survival curve after lipopolysaccharide (LPS) injection housed at 30°C and room temperature (n=10/group). **(L)** Circulating ketones, **(M)** rectal temperatures and **(N)** Kaplan–Meier survival curve after LPS with glucose supplementation (n=10/group). LPS dose was 10 mg/kg. Data are represented as mean ± SEM. *p < 0.05; **p<0.01; ***p < 0.001; ****p < 0.0001, log-rank (Mantel–Cox) test.

To determine whether hypoglycemia or hypothermia contribute to mortality during infection, we corrected hypoglycemia with glucose supplementation and prevented hypothermia by housing mice at thermoneutrality. Although thermoneutrality prevented hypothermia, it did not improve bradycardia, and paradoxically increased mortality (**Fig. 2I-K**). As expected, glucose supplementation significantly suppressed ketogenesis (**Fig. 2L**), but failed to improve hypothermia or survival (**Fig. 2M, N**). Together, these findings demonstrate that infection induces a conserved adipose lipolytic response associated with increased circulating NEFAs, and that hypothermia and hypoglycemia are correlated features rather than causal determinants of mortality.

### White adipose tissue lipolysis is necessary for tolerance to infection

To directly test whether adipose tissue lipolysis is required for tolerance to infection, we generated mice with adipose-specific deletion of ATGL (ATGL^AKO^), the rate-limiting enzyme for triglyceride hydrolysis(18). ATGL deletion was efficient in WAT and brown adipose tissue (BAT), with preserved expression in the liver (**Fig. 3A** and **Fig. S3A**). During CLP-induced sepsis, the increase in circulating NEFAs observed in control (ATGL^Flox^) mice was completely abolished in ATGL^AKO^ mice (**Fig. 3B**), and this response was similarly absent during LPS-induced inflammation (**Fig. 3C**).

**Figure 3:**
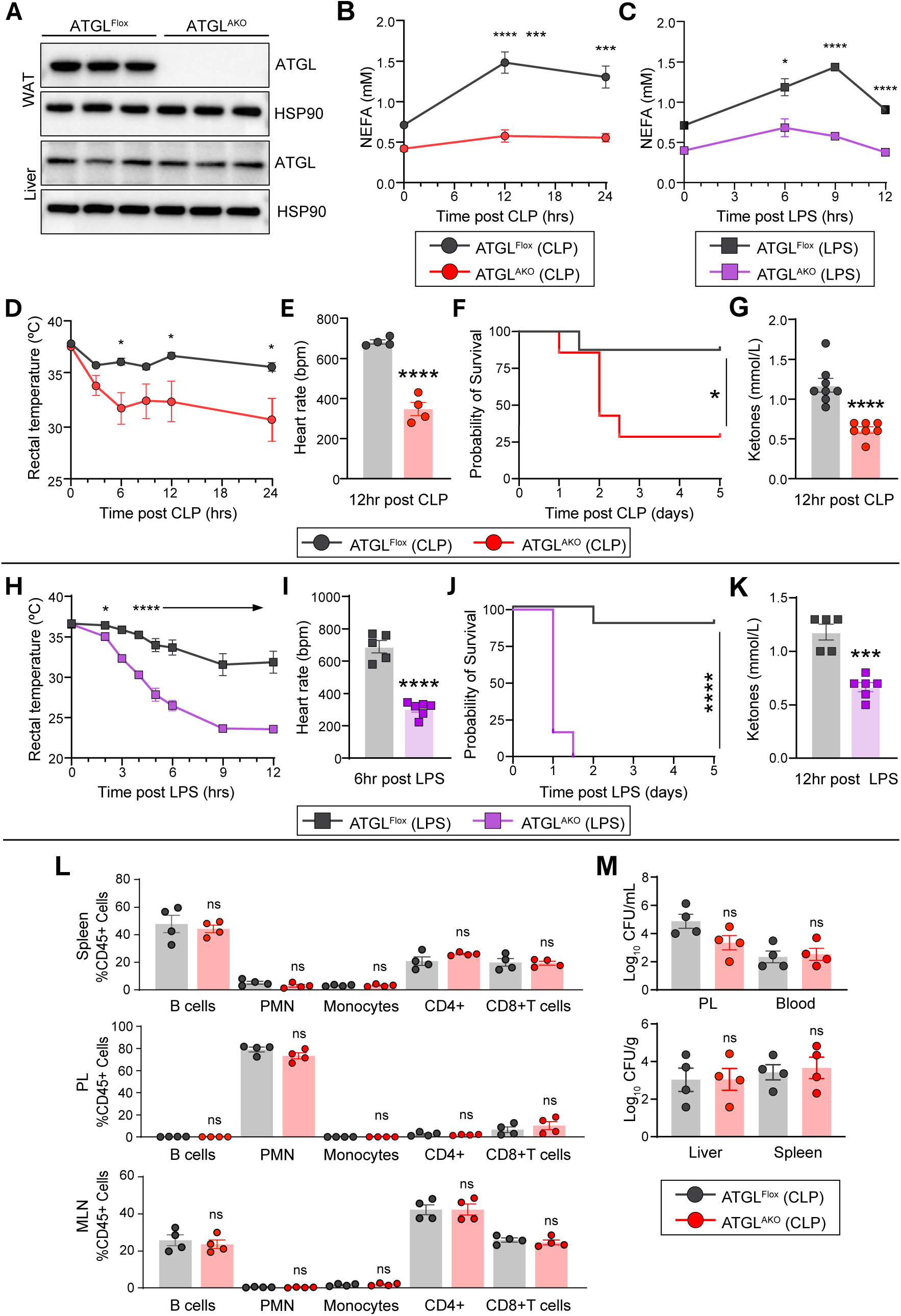
White adipose tissue lipolysis is necessary for tolerance to infection. **(A)** Immunoblot of ATGL expression in WAT and liver (n=3/group). Circulating **(B)** non-esterified fatty acids (NEFAs) after cecal ligation and puncture (CLP) (n=7-8/group). **(C)** Circulating NEFAs after lipopolysaccharide (LPS) (n=5/group). **(D)** Rectal temperatures (n=7-8/group), **(E)** heart rates (n=4/group), **(F)** Kaplan–Meier survival and **(G)** ketones after CLP (n=7-8/group). **(H)** Rectal temperatures, **(I)** heart rates and **(J)** Kaplan–Meier survival curve and **(K)** ketones after LPS (n=6-8/group). Immunophenotying of **(L)** spleen, peritoneal lavage (PL), mesenteric lymph node (MLN) 24 h after CLP (n=4/group). **(M)** Bacteria cultured from peritoneal lavage (PL), blood, liver, and spleen 24 h after CLP (n=4/group). Data are represented as mean ± SEM. *p < 0.05; **p<0.01; ***p < 0.001; ****p < 0.0001, log-rank (Mantel–Cox) test. For (C-K), LPS dose was 5 mg/kg. For CLP, n=5/group. For LPS, n=6/group. CFU, colony forming unit.

Loss of adipose lipolysis resulted in marked autonomic instability during infection. Compared to controls, ATGL^AKO^ mice developed more severe hypothermia and bradycardia, and exhibited significantly increased mortality following CLP (**Fig. 3D-F**). Although hypoglycemia was comparable between the two groups (**Fig. S3B**), ketone production was markedly attenuated in ATGL^AKO^ mice, consistent with impaired availability of adipose-derived fatty acids (**Fig. 3G**). Similar defects in thermoregulation, heart rate, survival, and ketogenesis were observed following LPS challenge, with comparable hypoglycemia between genotypes (**Fig. 3H-K** and **Fig. S3C**).

Despite these profound physiologic differences, inflammatory and immune responses were preserved in ATGL^AKO^ mice. Circulating levels of TNFα, IL-10, and IL-6 were comparable between genotypes in both CLP and LPS models (**Fig. S3D, E**). In the CLP model, major immune cell populations, including B cells, neutrophils (PMN), monocytes, and CD4^+^ and CD8^+^ T cells, were unchanged between ATGL^AKO^ and control mice across spleen, peritoneal lavage fluid, and mesenteric lymph nodes (**Fig. 3L** and **Fig. S3F**). Consistent with this, bacterial burden was comparable between ATGL^AKO^ and control mice in peritoneal lavage fluid, blood, liver and spleen (**Fig. 3M**). Collectively, these results indicate that loss of adipose ATGL impairs physiologic tolerance to infection without altering pathogen control or immune activation.

Because ATGL^AKO^ mice were generated using adiponectin-Cre, which targets both WAT and BAT, we next examined whether impaired tolerance to infection reflected loss of lipolysis in specific adipose tissues. In contrast to adiponectin-Cre-mediated deletion, BAT-specific deletion of ATGL using UCP1-Cre (ATGL^BKO^) did not impair circulating NEFAs, body temperature, heart rate, or survival compared to control mice following LPS challenge (**Fig. S3G-K**). Together, these findings demonstrate that lipolysis in WAT, but not BAT, is required to maintain autonomic stability and promote metabolic tolerance during infection.

### Infection-induced adipose lipolysis is redundantly regulated

WAT lipolysis can be activated by multiple upstream inputs, including adrenergic signaling, cytokine-mediated inflammatory pathways, and Toll-like receptor (TLR)-dependent signaling (15,21,22). To determine whether infection-induced lipolysis depends on any single canonical pathway, we disrupted key upstream regulators. Adipose-specific deletion of the β3-adrenergic receptor (ADRB3^AKO^) effectively impaired adrenergic signaling in WAT, as evidenced by reduced *Adrb3* expression and loss of LPS-induced phosphorylation of PKA substrates and HSL (**Fig. S4A, B**). However, compared to control mice, ADRB3^AKO^ mice exhibited intact NEFA release and no differences in body temperature, glucose, ketones or survival following CLP (**Fig. S4C-G**). Similar results were observed following LPS challenge (**Fig. S4H-L**). Pharmacologic inhibition of ERK1/2 and STAT3 signaling using the MEK inhibitor trametinib and the JAK inhibitor ruxolitinib, both FDA-approved agents, suppressed LPS-induced phosphorylation of ERK1/2 and STAT3, respectively, in WAT (**Fig. S4M, N**). However, these interventions did not attenuate the increase in circulating NEFAs (**Fig. S4O, P**). Together, these findings suggest that infection-induced adipose lipolysis does not depend on a single upstream signaling pathway but instead reflects a convergent and redundantly organized regulatory program that ensures lipid mobilization during systemic infection.

### Circulating fatty acids mediate lipolysis-dependent tolerance to infection

WAT lipolysis liberates two principal metabolites into circulation: NEFAs and glycerol (19). To determine which of these mediates lipolysis-dependent tolerance to infection, we selectively restored each metabolite in ATGL^AKO^ mice. To increase circulating NEFAs independently of WAT lipolysis, we administered intralipid together with heparin, providing triglyceride substrate and promoting lipoprotein lipase-dependent NEFA release, respectively (**Fig. S5A**).

In ATGL^AKO^ mice subjected to LPS-induced inflammation, intralipid increased circulating NEFAs and ketones to levels comparable to, or exceeding, those observed in control, ATGL^Flox^ mice (**Fig. 4A, B**). Restoring circulating NEFAs was sufficient to rescue metabolic tolerance, fully normalizing hypothermia and bradycardia to levels observed in control mice, and significantly improving survival compared to vehicle-treated ATGL^AKO^ mice (**Fig. 4C-E**). Intralipid treatment did not affect LPS-induced hypoglycemia (**Fig. 4F**). In contrast, glycerol supplementation increased circulating glycerol levels (**Fig. S5B**) but failed to improve body hypothermia, bradycardia, or survival in ATGL^AKO^ mice (**Fig. S5C-E**). Together, these findings demonstrate that circulating NEFAs, but not glycerol, mediate lipolysis-dependent tolerance to infection.

**Figure 4:**
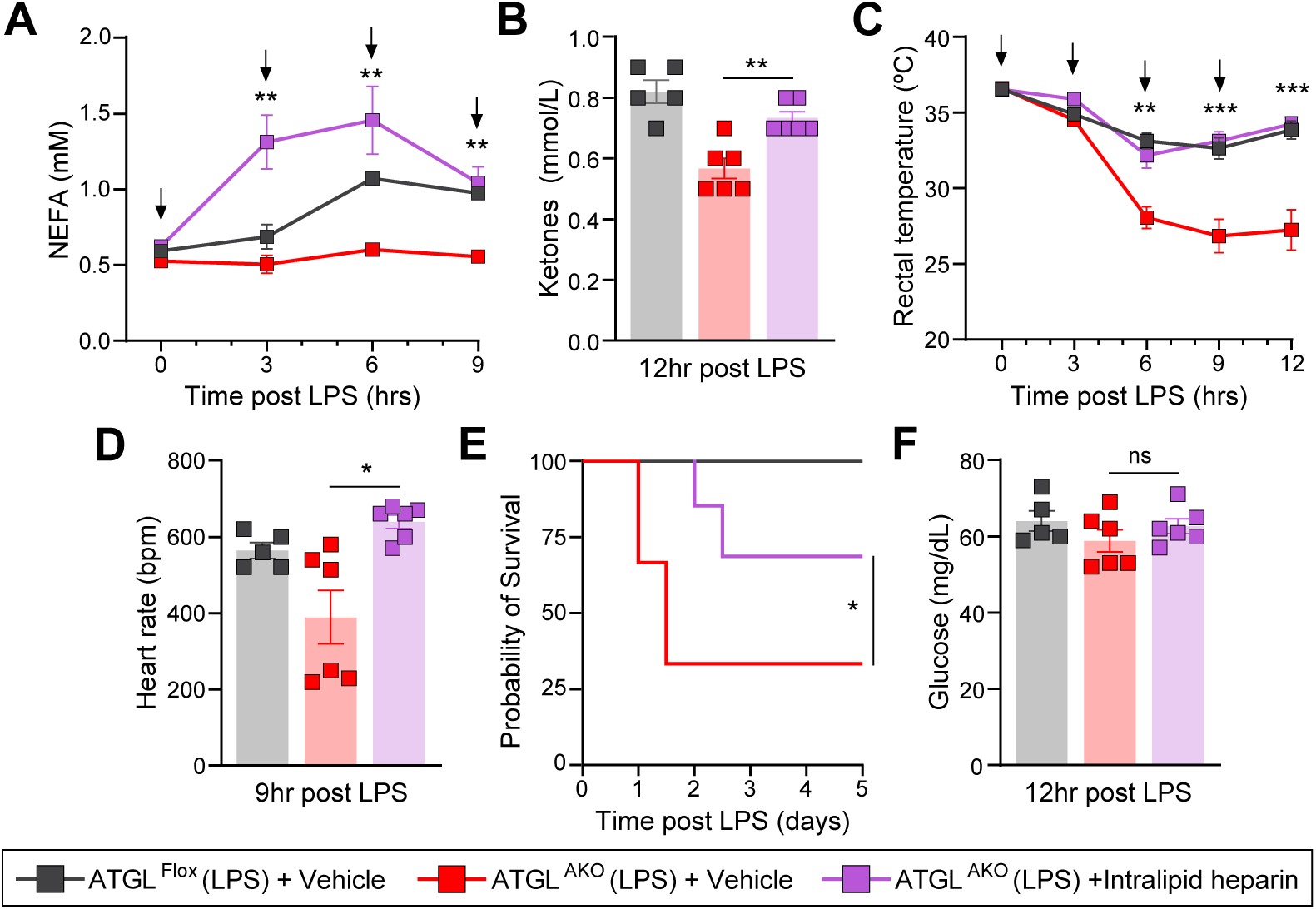
Circulating fatty acids mediate lipolysis-dependent tolerance to infection. **(A)** Circulating non-esterified fatty acids (NEFAs), **(B)** ketones, and **(C)** rectal temperatures, **(D)** heart rates, **(E)** Kaplan–Meier survival curve, and **(F)** glucose after liposaccharide (LPS) with intralipid heparin supplementation. The arrows indicate the time of intralipid heparin supplementation. LPS dose was 2.5 mg/kg and n=5-6/group. Data are represented as mean ± SEM. *p < 0.05; **p<0.01; ***p < 0.001; ****p < 0.0001, log-rank (Mantel–Cox) test.

### Mitochondrial fatty acid utilization in extrahepatic tissues promotes tolerance to infection

Our findings raise the question of where circulating fatty acids are utilized during infection and how they contribute to metabolic tolerance. The liver is a major sink for NEFAs released from adipose tissue during lipolysis, where these fatty acids undergo mitochondrial β-oxidation to support various metabolic processes such as ketogenesis (23). To test whether hepatic utilization of these lipids is required for tolerance to infection, we selectively disrupted mitochondrial fatty acid oxidation in the liver. CPT1a is the liver-enriched member of the carnitine palmitoyltransferase 1 family that catalyzes the rate-limiting step in mitochondrial import of long-chain fatty acids (LCFA) (24). We selectively deleted *Cpt1a* in hepatocytes using *Cpt1a floxed* mice and adeno-associated virus (AAV) expressing Cre recombinase under the control of the hepatocyte-specific TBG promoter (CPT1a^LKO^). Control mice were injected with AAV expressing TBG-GFP (CPT1a^Flox^). CPT1a^LKO^ mice showed efficient hepatic deletion and as expected, displayed significantly reduced circulating ketones in response to LPS challenge (**Fig. S6A, B**). Despite, this, CPT1a^LKO^ mice did not exhibit increased hypothermia, bradycardia, hypoglycemia or mortality following LPS challenge compared to control mice (**Fig. S6C-F**).

To exclude compensation by alternative CPT1 paralogs, we similarly deleted *Cpt2* in hepatocytes by administering AAV-TBG-Cre to CPT2^Flox^ mice (CPT2^LKO^). In contrast to CPT1, CPT2 lacks paralogs and is required for the final step of LCFA mitochondrial import (25). CPT2^LKO^ mice exhibited reduced hepatic *Cpt2* expression and, similar to CPT1a^LKO^ mice, had decreased circulating ketones following LPS challenge (**Fig. S6G, H)**. However, CPT2^LKO^ mice did not exhibit increased hypothermia, bradycardia, hypoglycemia, or mortality compared to control mice (**Fig. S6I-L**). Together, these data demonstrate that hepatic mitochondrial fatty acid oxidation is not required for metabolic tolerance to infection.

Since NEFAs liberated during lipolysis can undergo CPT1-dependent oxidation in extrahepatic tissues, we next inhibited CPT1 systemically using etomoxir (26). Etomoxir treatment reduced circulating ketones and glucose, and increased NEFA levels during CLP, confirming effective CPT1 blockade (**Fig. 5A-C**). Compared to vehicle, etomoxir treatment significantly worsened infection-induced hypothermia, bradycardia, and mortality (**Fig. 5D-F**). Similar effects of etomoxir treatment were observed following LPS challenge (**Fig. 5G-K**). Because etomoxir has been reported to exert off-target mitochondrial effects (27,28), we next tested whether the observed phenotype reflected inhibition of CPT1-dependent fatty acid transport rather than global disruption of mitochondrial function. Supplementation with medium-chain triglycerides (MCT oil), which bypass CPT1-dependent mitochondrial import, restored ketone production in etomoxir-treated mice challenged with LPS, whereas long-chain triglycerides (corn oil) did not (**Fig. S6M**). Together, these findings demonstrate that CPT1-dependent mitochondrial utilization of fatty acids is required to maintain tolerance during infection and is mediated by extrahepatic, rather than hepatic, fatty acid oxidation.

**Figure 5:**
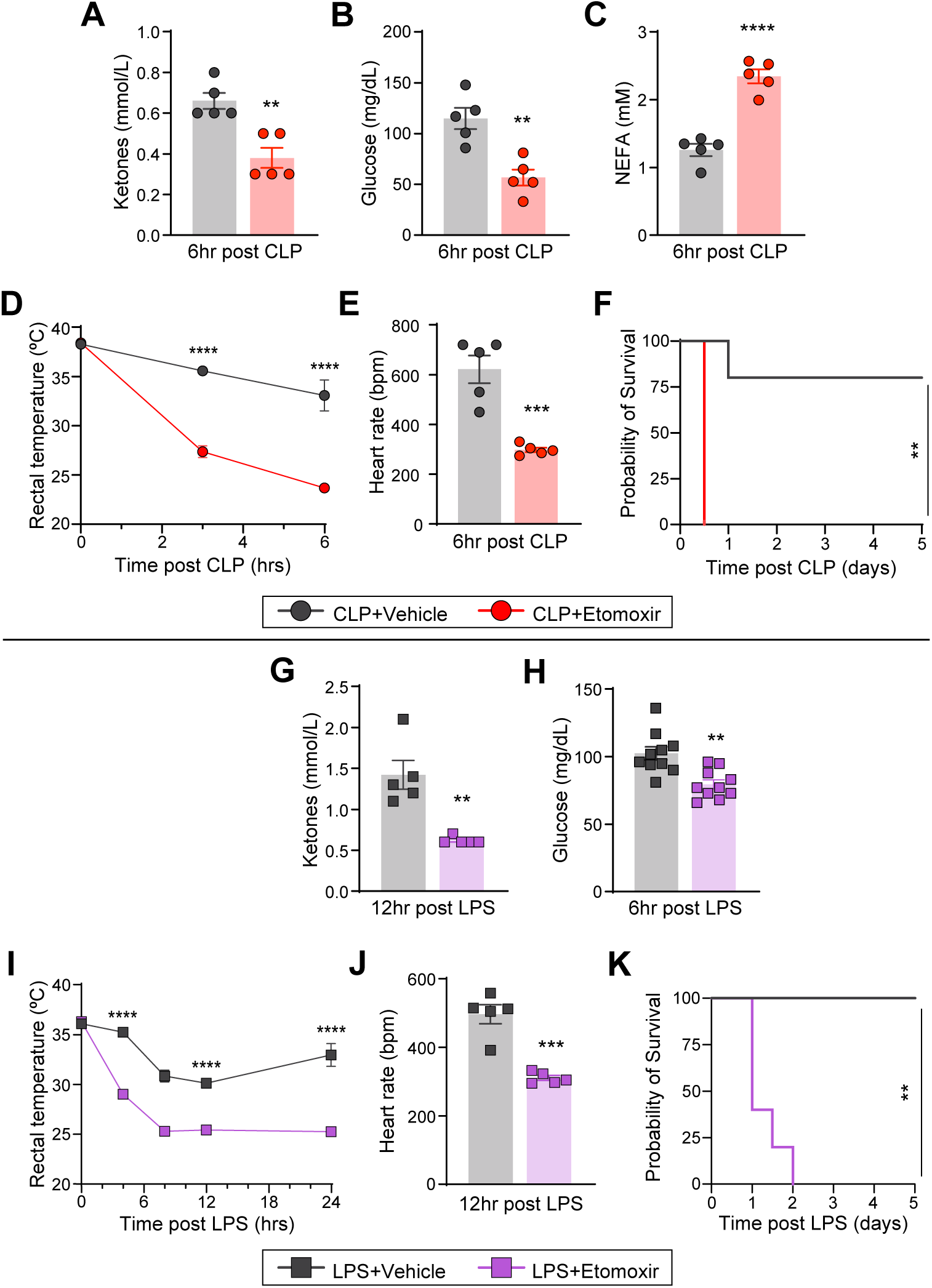
Extrahepatic fatty acid oxidation is required for tolerance to infection. Circulating **(A)** ketones, **(B)** glucose, **(C)** non-esterified fatty acids (NEFAs), **(D)** rectal temperatures, **(E)** heart rates, and **(F)** Kaplan–Meier survival curve after cecal ligation and puncture (CLP) and etomoxir. (n=5/group). Circulating **(G)** ketones, **(H)** glucose, **(I)** rectal temperatures, **(J)** heart rates, **(K)** Kaplan–Meier survival curve after LPS and etomoxir. For (G-K), LPS dose was 5 mg/kg and n=5-10/group. Data are represented as mean ± SEM. *p < 0.05; **p < 0.01; ***p < 0.001; ****p < 0.0001, log-rank (Mantel–Cox) test.

### Induction of adipose lipolysis promotes tolerance to inflammation

Although β3-adrenergic signaling in adipose tissue was not required for tolerance to infection (**Fig. S4**), we next tested whether pharmacologic induction of lipolysis was sufficient to promote tolerance. Treatment with the β3-adrenergic agonist CL-316,243 (CL3) during LPS challenge rapidly increased circulating NEFA and ketone levels without altering glucose (**Fig. S7A-C**). CL3 treatment protected against LPS-induced hypothermia and bradycardia, and significantly improved survival (**Fig. 6A-C**), indicating that induction of lipolysis is sufficient to enhance tolerance to inflammation. Since LPS challenge is associated with reduced energy expenditure (5), we performed indirect calorimetry to assess the metabolic effects of CL3 in the context of sterile inflammation. CL3-treated mice exhibited increased energy expenditure and respiratory exchange ratio (**Fig. 6D, E**). To confirm that these effects were mediated through β3-adrenergic signaling in adipose tissue, we treated mice lacking adipose tissue β3-adrenergic receptors (ADRB3^AKO^) and control littermates (ADRB3^Flox^) with CL3 during LPS challenge. In control ADRB3^Flox^ mice, CL3 increased circulating NEFA and ketone levels, with comparable glucose levels and protected against LPS-induced hypothermia, bradycardia, and mortality, compared to vehicle treatment. In contrast, these metabolic and physiologic effects were completely absent in ADRB3^AKO^ mice treated with CL3 (**Fig. 6F-H** and **Fig. S7D-F**), confirming that the protective effects of CL3 require β3-adrenergic signaling in adipose tissue.

**Figure 6:**
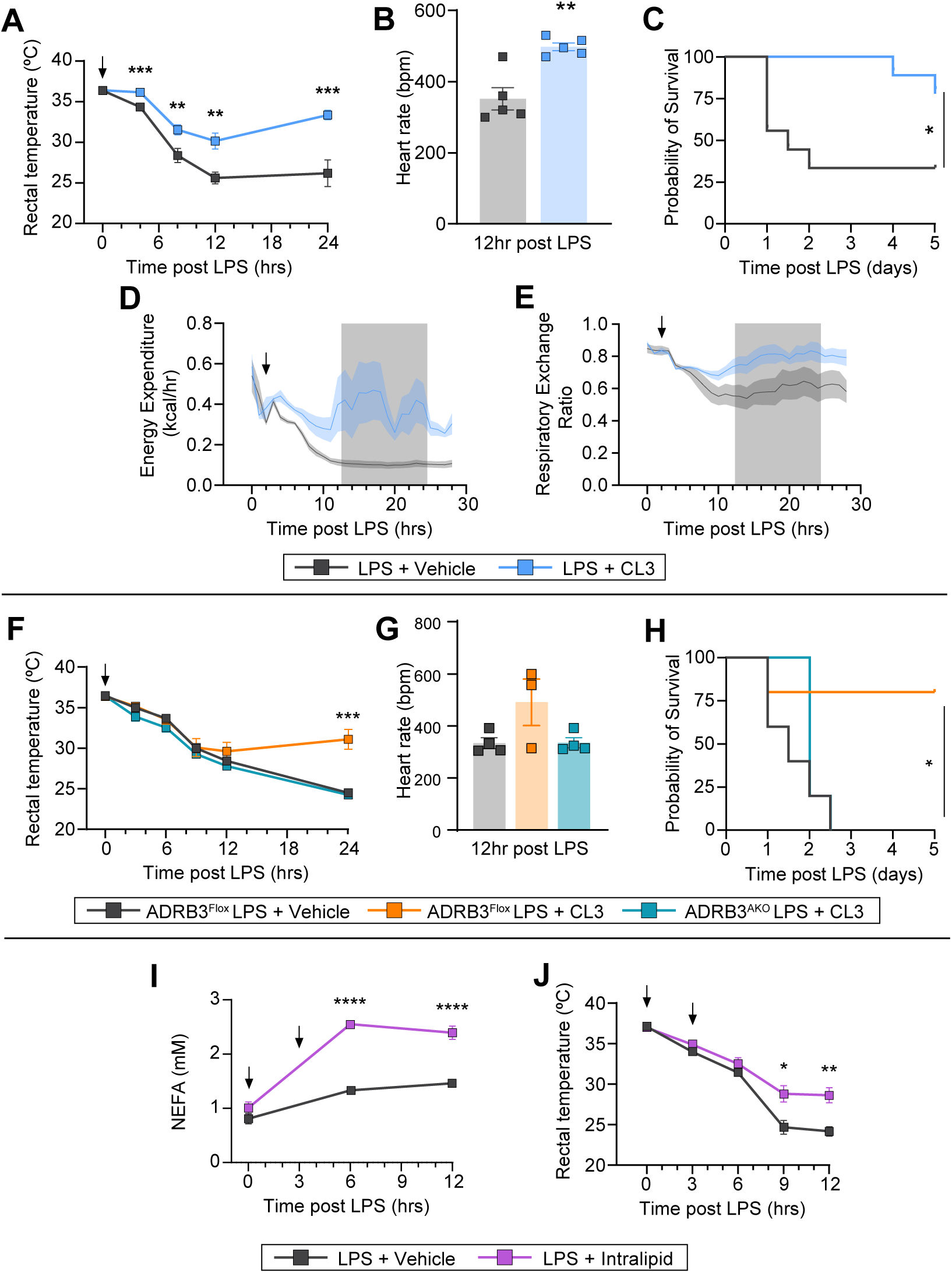
Inducing lipolysis promotes tolerance to inflammation. **(A)** Rectal temperatures (n=9/group), **(B)** heart rates (n=5/group), **(C)** Kaplan–Meier survival curve (n=9/group), **(D)** energy expenditure (n=4/group), and **(E)** respiratory exchange ratio (n=4/group) after lipopolysaccharide (LPS) and CL316,243 (CL3). **(F)** Rectal temperatures, **(G)** heart rates, and **(H)** Kaplan–Meier survival curve after LPS and CL3 (n=4-6/grourp). **(I)** Circulating non-esterified fatty acids (NEFAs), **(J)** rectal temperatures after LPS with intralipid supplementation (n=6-7/group). LPS dose was 10 mg/kg. Arrows indicate the time of CL3 administration and intralipid supplementation. Data are represented as mean ± SEM. *p < 0.05; **p < 0.01; ***p < 0.001; ****p < 0.0001, log-rank (Mantel–Cox) test.

To further test whether increasing circulating fatty acids is sufficient to promote tolerance, we supplemented mice with intralipid during LPS challenge. Intralipid treatment rapidly and durably increased circulating NEFA levels and protected against LPS-induced hypothermia (**Fig. 6I, J**). Together, these findings demonstrate that both induction of lipolysis and exogenous supplementation of circulating fatty acids are sufficient to promote tolerance to inflammation.

### β3-adrenergic receptor agonist exposure is associated with improved survival in patients with sepsis

Mirabegron and vibegron are FDA-approved β3-adrenergic receptor agonists indicated for overactive bladder in adults (29,30). During clinical trials, plasma NEFA levels were significantly higher after starting mirabegron treatment compared to before (31,32). To determine whether β3 agonist exposure is associated with clinical benefit in sepsis, we identified patients admitted with sepsis between 2022 and 2024 who received mirabegron within the first 24 hours of hospitalization using the Epic Cosmos EHR database (33). Patients were propensity score matched on clinical and encounter-level covariates (**Tables 1** and **S1**). Exposure to a β3 agonist was associated with significantly reduced risk of mortality or hospice discharge (OR 0.66; 95% CI, 0.60-0.72; **Fig. 7A**). This association was consistent across prespecified demographic subgroups, including male, female, White, and Black patients. A dose-response analysis further revealed that patients who received 50 mg had lower risk of mortality or hospice discharge compared to those who received 25 mg (OR 0.66; 95% CI, 0.60-0.72; **Fig. 7B**). Together, these findings demonstrate that β3 agonist exposure is associated with a dose-dependent reduction in mortality among patients with sepsis.

**Figure 7:**
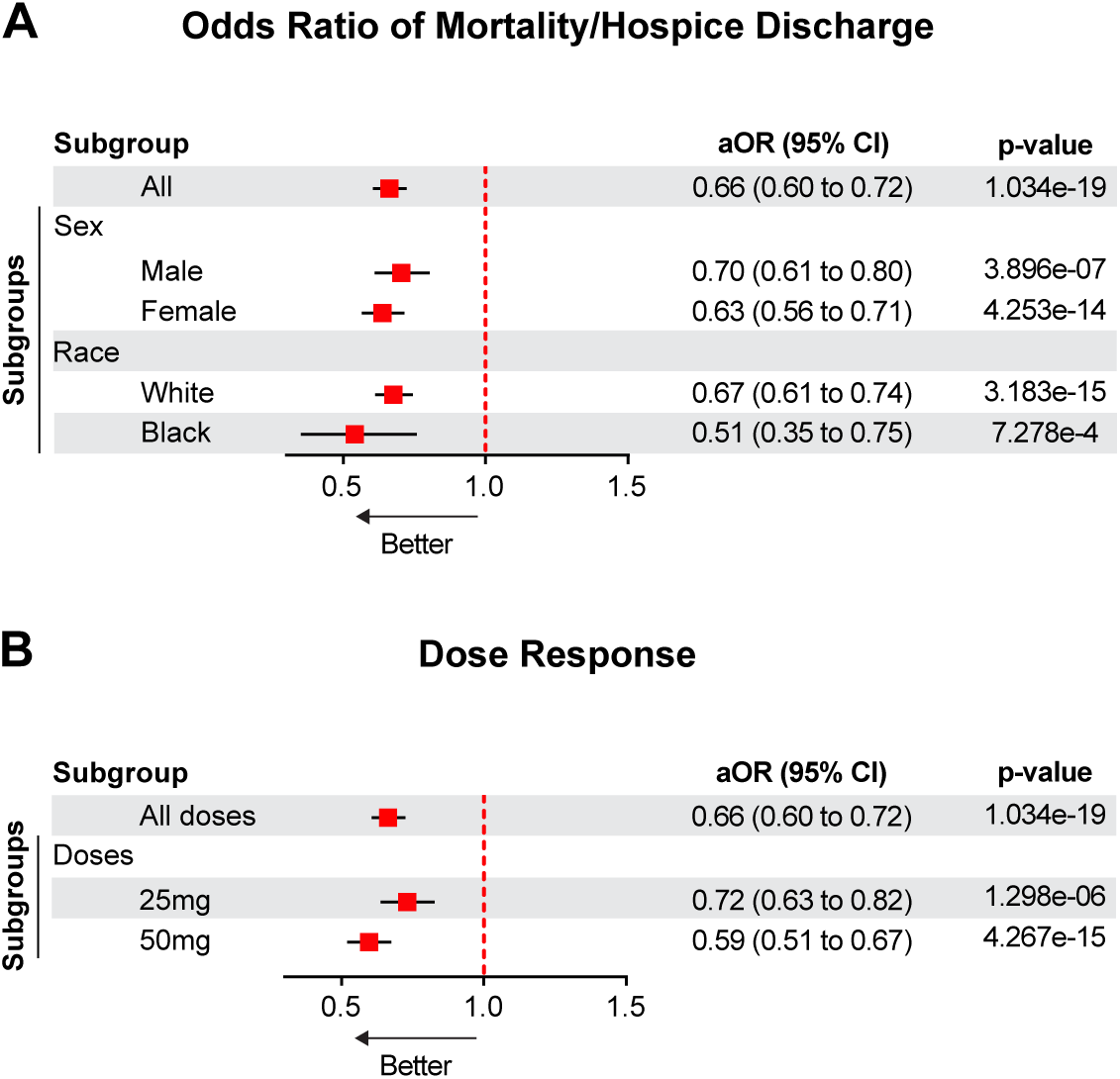
Exposure to a β3 agonist improved survival in patients with sepsis. **(A)** Forest plot of odds ratios for mortality or hospice discharge associated with β3 agonist exposure. **(B)** Forest plot of odds ratios for mortality or hospice discharge by β3 agonist dose.

## DISCUSSION

In this study, we identify adipose tissue lipolysis as a critical metabolic mechanism that promotes tolerance to infection. Using genetic, pharmacologic, and nutritional approaches, we show that mobilization of fatty acids from adipose tissue is required to maintain autonomic stability and survival during endotoxemia and polymicrobial sepsis. Loss of adipocyte lipolysis impaired circulating NEFA availability and resulted in hypothermia, bradycardia, and mortality despite preserved inflammatory responses and pathogen burden. Restoration of circulating fatty acids rescued these phenotypes, whereas glucose or glycerol supplementation did not, indicating that lipid-derived fuels rather than carbohydrate substrates are central to this adaptive response. Mechanistically, our data suggests extrahepatic mitochondrial fatty acid oxidation as an important downstream pathway supporting tolerance during infection. Together with our human data linking circulating NEFA levels and pharmacologic stimulation of lipolysis to improved sepsis outcomes, these findings identify adipose lipid mobilization as a key metabolic component of host tolerance during infection.

Our data suggests that mitochondrial fatty acid oxidation is an important component of promoting tolerance during infection. Pharmacologic inhibition of CPT1-mediated fatty acid oxidation with etomoxir phenocopied adipose ATGL deficiency, producing hypothermia, bradycardia, and mortality in both endotoxemia and polymicrobial sepsis models. Etomoxir can have off-target effects, including inhibition of mitochondrial complex I, CoA sequestration, and global disruption of mitochondrial metabolism (26,27). However, although etomoxir-treated mice failed to generate ketones in response to long-chain triglycerides (corn oil), ketogenesis was robustly restored when animals were supplemented with medium-chain triglycerides (MCT oil), which bypass CPT1-dependent mitochondrial import (34). These findings argue against global mitochondrial dysfunction and instead suggest a requirement for CPT1-dependent mitochondrial utilization of adipose-derived fatty acids during infection.

A key unresolved question raised by our findings is where circulating fatty acids are utilized during infection and how they promote tolerance. The liver is often presumed to be the dominant sink for adipose-derived fatty acids during systemic stress, in part because hepatic fatty acid oxidation drives ketone production during fasting and infection (34,35). Surprisingly, however, hepatocyte-specific deletion of CPT1a or CPT2 did not recapitulate the ATGL-deficient phenotype, arguing against a primary requirement for hepatocyte long-chain fatty acid oxidation or ketone production during infection. This interpretation is consistent with emerging genetic evidence suggesting that hepatic ketogenesis is dispensable for survival and physiologic responses during endotoxemia (36–38). Together, these observations raise the possibility that extrahepatic tissues, including cardiovascular, skeletal muscle, neural, or immune compartments, oxidize adipose-derived fatty acids to sustain key features of physiological tolerance such as thermoregulation and heart rate. In addition to serving as substrates for fuel, circulating fatty acids may also function as signaling molecules, for example as ligands for lipid-sensing nuclear receptors such as PPARs, suggesting that lipid mobilization may coordinate tolerance through both metabolic and transcriptional programs (23,39,40).

Although adipose lipolysis is required to establish tolerance to infection, glucose supplementation is not. In our models, glucose supplementation neither improved nor worsened hypothermia, bradycardia, or survival following endotoxemia or polymicrobial sepsis. However, prior work reported that glucose administration worsened mortality during endotoxemia and *Listeria* infection, suggesting that glucose itself may be detrimental (36). Notably, in that study glucose was administered in water while control animals received PBS gavage. In our experiments glucose was dissolved in PBS with a matched PBS-vehicle control. When we repeated the experiment with glucose dissolved in water and PBS used as the vehicle, we similarly observed worsened mortality following LPS, consistent with Wang et al (**Fig. S8**) (36). These observations suggest that differences in experimental formulation or vehicle composition may underlie the physiological effects attributed to glucose supplementation. Importantly, numerous earlier studies also reported that glucose supplementation neither protected nor worsened survival in LPS-challenged mice (41,42). Clarifying the impact of glucose administration during infection is clinically important, as patients with sepsis routinely receive glucose through intravenous fluids and enteral nutrition containing glucose polymers such as maltodextrin.

Our findings indicate that infection-induced adipose lipolysis does not depend on any single upstream signaling pathway. Prior studies have shown that inflammatory stimuli can activate lipolytic signaling in adipocytes through multiple mechanisms. Experimental endotoxemia in humans induces phosphorylation of HSL in adipose tissue, supporting the idea that systemic inflammation directly engages adipocyte lipolysis *in vivo* (13). Mechanistic studies further demonstrate that TLR4 activation can stimulate ERK signaling, and that inflammatory cytokines can activate MAPK and JAK/STAT pathways to promote lipolysis in adipocytes (15,16,43). However, many of these observations derive from *in vitro* or cell-autonomous systems and do not fully capture the integrated physiology of infection, in which multiple inflammatory and neuroendocrine signals are activated simultaneously. In contrast, our *in vivo* genetic and pharmacologic studies show that disruption of individual pathways, including β3-adrenergic signaling, ERK activation, or JAK/STAT signaling, does not impair infection-induced lipid mobilization. Together, these findings support a model in which adipose lipolysis during systemic infection is governed by convergent and partially redundant regulatory inputs that ensure reliable fatty acid availability during inflammatory stress.

An additional implication of our findings relates to the so-called “obesity paradox” observed in sepsis and other critical illnesses, in which patients with higher body mass index exhibit improved survival compared with lean individuals (44–46). Although the mechanisms underlying this phenomenon remain debated, one possibility is that greater adipose energy reserves provide a metabolic advantage during severe systemic inflammation. Consistent with this, sepsis is a leading cause of premature death in individuals with lipodystrophy, especially in Congenital Generalized Lipodystrophy (CGL) patients (47). Experimental studies in murine models have produced mixed results, with some reports demonstrating improved survival in obese mice subjected to polymicrobial sepsis, whereas others show worsened or unchanged outcomes, highlighting the complexity of metabolic influences on host responses to infection (48,49). Our findings raise the possibility that the capacity to mobilize adipose-derived fatty acids during infection may represent one metabolic mechanism linking adiposity to improved tolerance in severe infection.

Our findings have translational implications. Circulating NEFAs are known to increase in patients with sepsis as a consequence of infection-induced lipolysis (50,51). Consistent with these observations, we found that circulating NEFA levels were elevated in patients with sepsis and that higher NEFAs were independently associated with improved survival and lower clinical severity. These findings suggest that lipid availability reflects physiologic tolerance rather than disease progression and parallel our murine data supporting an adaptive role for infection-induced lipolysis. Consistent with this interpretation, analysis of nationwide real-world EHR data demonstrated that use of β3-adrenergic agonists, including mirabegron, was associated with lower risk of inpatient mortality among patients admitted with sepsis. This agrees with our mechanistic findings implicating adipose lipolysis in host survival. However, this EHR-based data is not without limitations. Because β3-adrenergic agonists are prescribed for overactive bladder, the population receiving them is systematically healthier/functionally independent than those who are not. Despite propensity matching, residual confounding can’t be fully excluded. However the dose-response relationship observed is more consistent with a pharmacologic effect than with a healthy-user bias. These observations raise important questions regarding the metabolic management of infected patients, including the composition of nutritional support in the intensive care unit. Our findings suggest that preservation of fatty acid availability during infection may be beneficial and warrant further mechanistic and clinical investigation.

## MATERIALS AND METHODS

### STUDY DESIGN

The objective of this study was to define the role of adipose tissue lipolysis and circulating fatty acids in promoting tolerance to infection and to evaluate the translational relevance of these findings in human sepsis. We combined analyses of human clinical data with mechanistic studies in mouse models of infection and inflammation.

In human studies, plasma metabolomic profiling from the Early Acute Renal and Lung Injury (EARLI) cohort was analyzed to determine associations between circulating NEFA levels and sepsis severity and mortality. These findings were complemented by an independent retrospective cohort analysis using Epic Cosmos to evaluate whether exposure to β3-adrenergic receptor agonists within the first 24 hours of hospitalization was associated with improved outcomes among patients with sepsis. The primary outcome of the clinical cohort study was a composite of in-hospital mortality or discharge to hospice.

In parallel, we used established mouse models of polymicrobial sepsis (cecal ligation and puncture, CLP) and endotoxemia (lipopolysaccharide, LPS) to define the role of adipose lipolysis in infection. Genetic loss-of-function approaches included adipose-specific deletion of adipose triglyceride lipase (ATGL) and β3-adrenergic receptor (ADRB3), as well as tissue-specific deletion of enzymes involved in mitochondrial fatty acid oxidation. Pharmacologic interventions included β3-adrenergic agonists, inhibitors of signaling pathways regulating lipolysis, and lipid supplementation (Intralipid) strategies to modulate circulating fatty acid availability.

Physiologic outcomes included body temperature, heart rate, glucose, circulating metabolites, and survival. Immune responses were assessed by flow cytometry and cytokine analyses, and pathogen burden was quantified by colony-forming assays. Investigators were not blinded to genotype during experiments. Animals were randomly assigned to experimental groups, and sample sizes were selected based on prior studies and pilot data to ensure adequate power to detect biologically meaningful differences. For survival studies, predefined humane endpoints were applied according to institutional animal care guidelines. Statistical methods and numbers of biological replicates are described in the relevant figure legends and Statistical Analysis section.

### Mice

All experiments were conducted on C57BL6/J male mice, which were 7-8 weeks of age and housed in a 12-h light/12-h dark cycle. ATGL^AKO^ mice (generated by crossing *Atgl^fl/fl^* mice with *Adipoq-Cre* mice) were a gift from Drs. Evan Rosen and Barbara Kahn. ADRB3^AKO^ mice were generated by crossing *Adrb3*^fl/fl^ mice and *Adipoq-Cre* mice. ATGL^BKO^ mice were generated by crossing *Atgl*^fl/fl^ mice and *Ucp1-Cre* mice (Jackson Labs # 024670). *Cpt1a^fl/fl^* and *Cpt2^fl/fl^* mice were gifts from Dr. Jeffrey Friedman and Dr. Michael J. Wolfgang, respectively. 5-week-old *Cpt1a^fl/fl^* and *Cpt2^fl/fl^* mice were retro-orbitally injected with 2 × 10^11^ viral genomes(vg) of AAV8 expressing Cre recombinase or GFP under the control of hepatocyte-specific TBG promoter (AAV8-*TBG*-*Cre* and AAV8-*TBG*-*GFP*). For making the AAVs, pAAV8 vectors were transfected in cultured CRL3022 cells (HEK293S GnTI) followed by purification. Experiments were performed 2-3 weeks after injection of the purified virus.

Mice were intraperitoneally injected with the mentioned dose of LPS derived from *Escherichia coli* 055:B5 (Sigma-Aldrich) diluted with PBS. While controlled within each experiment, LPS doses ranged from 2.5 to 10 mg/kg based on mouse strain specific susceptibility and differences in the potency of each LPS batch/lot, with specific doses contained in the main text.

For induction of polymicrobial sepsis, mice were subjected to cecal ligation and puncture (CLP) as described (52,53). Mice were anesthetized by isoflurane inhalation, and under aseptic conditions, a one-centimeter incision was made in the abdomen to externalize the cecum and 50% (precisely 1.2 cm) was ligated below the ileocecal valve. This was followed by puncturing the cecum with a 27-gauge needle for developing mild sepsis. During the procedure, some cecal content was expelled using sterile forceps. The abdominal musculature and skin were closed with simple running sutures and metallic clips, respectively. Buprenorphine at 0.5 mg/kg was administered after CLP according to the IACUC protocol. Sham-operated mice served as controls. Sham procedures consisted of laparotomy with cecal exposure without ligation or puncture.

For survival studies, mice were single-housed and acclimatized for 2 days prior to LPS injection or CLP surgery. According to the IACU protocol, mice were euthanized if their core temperature fell to ≤23°C or if they lost their righting reflex (survival studies).

Mice were gavaged with vehicle or the equivalent of one kilocalorie glucose twice daily after LPS injection. Trametinib (8 mg/kg) and Ruxolitinib (50 mg/kg) were administered by oral gavage 90 minutes after LPS injection in 200 µL of 0.5% (v/v) methylcellulose. A bolus of 20% intralipid and heparin (10 U per injection) was administered intraperitoneally after LPS injection. The timing of administration is indicated by arrows in respective figures. Glycerol (2.0 g/kg) was administered intraperitoneally 3 h after LPS injection. Etomoxir (30 mg/kg) was injected intraperitoneally every 12 h after LPS or CLP. CL316,243 (CL3) was administered intraperitoneally (250 ug/kg) 30 minutes after LPS injection.

MouseOx® Plus (Starr Life Sciences) was used for monitoring the heart rate on unanesthetized, freely moving mice after LPS injection or CLP as previously described (54).

Lubricated rectal thermometer (Digi-Sense) was used for measuring core body temperature. For thermoneutrality experiments, mice were housed at 30°C. Following LPS administration, animals were immediately transferred to temperature-controlled chambers maintained at 30°C for the duration of the study. Core body temperature was continuously measured by radiotelemetry using intra-abdominal implantable transmitters (DSI). Telemetry probes were surgically implanted into the peritoneal cavity under isoflurane anesthesia using aseptic technique, and mice were allowed to recover before experimentation. Temperature data were acquired using the DSI data acquisition system and analyzed.

For metabolic measurements, mice were individually housed in Promethion metabolic phenotyping cages (Sable Systems) under a 12 h light–dark cycle. Mice were acclimated for a minimum of 7 days before starting the experiment. Acclimation was confirmed by stabilization of measured metabolic parameters. Following acclimation, mice were intraperitoneally injected with LPS (10 mg/kg) at ZT2. 30 minutes after LPS administration, mice were intraperitoneally injected with CL3 (250 ug/kg). Metabolic parameters were continuously recorded from the time mice were placed into the metabolic cages up to 24 h after LPS injection. At the conclusion of the recording period, mice were removed from the cages and euthanized.

All animal experiments were conducted under protocols approved by UT Southwestern or Janelia Research Campus.

### Statistical Analysis

Statistical analyses were performed using GraphPad Prism 10. All graphs and data are presented as mean ± standard error of the mean. Statistical significance was determined using the unpaired two-tailed Student’s *t* test for single variables. A p value of < 0.05 was statistically significant and is presented as ^∗^ (p < 0.05), ^∗∗^ (p < 0.01), ^∗∗∗^ (p < 0.001), or ^∗∗∗∗^ (p < 0.0001). P values for mouse survival curves were calculated using Mantel-Cox or Gehan-Breslow-Wilcoxon test. The number of biological replicates used in each of the studies are indicated in the figure legends as “n=.”

#### List of Supplementary Materials

Materials and Methods Fig. S1 to S8

Table S1 References (55-57)

## Acknowledgments

We thank Dr. Barbara B. Kahn and Dr. Evan D. Rosen (Beth Israel Deaconess Medical Center) for generously providing Atgl^fl/fl^:Adipoq-Cre mice. Cpt1a^fl/fl^ mice were kindly provided by Dr. Jeffrey Friedman (The Rockefeller University), and Cpt2^fl/fl^ mice were generously provided by Dr. Michael J. Wolfgang (Johns Hopkins Medicine). We also thank the Microarray and Immune Phenotyping Core (UT Southwestern Medical Center) for assistance with multiplex cytokine assays. The authors thank Drs. Philipp Scherer, David Beckham, Lance Terada, and Ralph DeBerardinis for helpful discussions.

## Author contributions

S.J.P. and K.N.R. conceptualized the work. A.S., J.Y., S.P., S.X., T.G., S.K., S.H., N.A., A.S., R.S., L.G., L.K. performed experiments. A.S., S.M.M.A.R., N.A.L. analyzed data. A.S., S.H., S.R., A.M.T., D.G.T., C.C., M.B., X.R., A.I., L.J., V.N., J.K.E., H.W.S.D., E.J.S., C.S.C., M.A.M., J.H. performed data curation. S.J.P., K.N.R., A.S. wrote the manuscript. S.J.P. and K.N.R. acquired funding. S.J.P. and K.N.R. supervised the work.

## Data and materials availability

All data associated with this study are present in the paper or the Supplementary Materials.

## SUPPLEMENTARY MATERIALS

### Materials and Methods

#### Human Studies

##### Circulating non-esterified fatty acids in healthy and infected patients

Healthy and infected patients were enrolled from primary care clinics and the surgical ICUs at Weill Cornell Medicine and the University of Cincinnati. Patients over the age of 18 years old admitted with a documented infection were enrolled into the study. In addition to patients whom we were unable to consent, patients were excluded if they were immunosuppressed (either from medications or from an autoimmune disease) or were actively receiving chemotherapy within the past 6 months. Once enrolled, demographic information, clinical data, and blood draws of up to 12 ml were collected within the first 48 hours of admission. This study was approved by the Institutional Review Board at Weill Cornell Medicine (24–04027385) and the University of Cincinnati (2019–1098).

##### EARLI Study Cohort

We analyzed plasma samples from 200 septic patients enrolled in the Early Acute Renal and Lung Injury (EARLI) cohort between 2008 and 2016. EARLI is a prospective observational study conducted at two University of California, San Francisco-affiliated hospitals (UCSF Moffit-Long Medical Center and Zuckerberg San Francisco General Hospital) that enrolls critically ill adults at the time of ICU admission from the emergency department. Patients admitted primarily for isolated neurological, neurosurgical, or traumatic injuries are excluded from enrollment. Sepsis cases were identified through retrospective adjudication by physician reviewers who evaluated electronic health record data according to Sepsis-2 definitions (55). This adjudication process incorporated comprehensive clinical and microbiologic information and was performed with physician reviewers masked to biological profiling results.

Metabolomic profiling was performed on 150 μL citrated plasma samples using an untargeted approach by Metabolon as described for a prior study (56). In brief, three complementary analytical platforms were employed: (1) reverse-phase ultra performance liquid chromatography-tandem mass spectrometry (RP/UPLC-MS/MS) in positive ion mode, (2) RP/UPLC-MS/MS in negative ion mode, and (3) hydrophilic interaction liquid chromatography UPLC-MS/MS (HILIC/UPLC-MS/MS) in negative ion mode. Peak identification was conducted in 2017 using Metabolon’s proprietary spectral library, with standard quality control procedures and batch effect correction applied.

Six circulating non-esterified fatty acids (NEFAs) were identified: linoleate (18:2n6), myristate (14:0), oleate/vaccenate (18:1), palmitate (16:0), palmitoleate (16:1n7), and stearate (18:0).

Metabolite identifications were confirmed by matching Metabolon compound identifiers to biochemical annotations.

Individual NEFA values (area under the curve of the relative peak intensity) were log-transformed as log (NEFA + 1) to address right-skewed distributions while ensuring all transformed values remained non-negative. A composite NEFA score was then calculated for each subject as the arithmetic mean of the six log-transformed NEFA values.

The composite NEFA score was compared across Acute Physiology and Chronic Health Evaluation (APACHE II) severity tertiles, 60-day mortality status (survived vs. died), and vasopressor use at enrollment (yes vs. no). The Shapiro-Wilk test was used to assess normality within each group. For comparisons involving three groups (APACHE II tertiles), either ANOVA or the Kruskal-Wallis test was used depending on normality, with a pairwise Wilcoxon rank-sum test comparing the lowest and highest tertiles. For two-group comparisons (mortality, vasopressor use), either Student’s t-test or the Wilcoxon rank-sum test was applied based on the normality assessment.

Multivariable logistic regression was used to estimate the odds ratio (OR) for 60-day mortality per standard deviation (SD) increase in each individual log-transformed NEFA and the composite NEFA score, adjusted for age, sex, and diabetes. Kaplan-Meier survival curves were generated by stratifying patients into high and low composite NEFA groups based on the median value. The log-rank test was used to compare the unadjusted survival distributions between groups. A multivariable Cox proportional hazards model estimated the hazard ratio (HR) for time to death per SD increase in the composite NEFA score, adjusted for age, sex, and diabetes. Patients alive at the end of follow-up were censored at 60 days. All analyses were performed in R version 4.5.1.

##### Epic Cosmos

This retrospective cohort analysis was conducted using Epic Cosmos. Epic Cosmos is a real-world EHR-derived dataset created in collaboration with a community of health systems, aggregating de-duplicated patient records of more than 300 million patients.

We identified the hospitalizations between 2022 and 2024 where the patient was an adult (aged ≥ 18 years) and had sepsis. Sepsis was defined using the Angus criteria, requiring the presence of at least one ICD-10-CM diagnosis code consistent with infection and at least one ICD-10-CM diagnosis code indicating acute organ dysfunction during the same hospitalization. We only considered encounters where the length of stay was between 3 and 30 days to remove extreme outliers and reduce bias from atypical sepsis admissions. If a patient had multiple encounters satisfying the inclusion criteria, only the first encounter was considered for the analysis.

The exposure was defined as administration of a β3 agonist (Mirabegron or Vibegron) within first 24 hours from admission. The primary outcome was composite of mortality or hospice discharge at the end of the hospitalization. For the time to event analysis, time zero was defined as the start of hospitalization and patients were followed until discharge. We performed a propensity-score matched analysis. Propensity scores were estimated using a multivariable logistic regression model including predefined baseline covariates. We did 1:1 nearest neighbor matching without replacement, and covariate balance was evaluated using standardized mean differences (SMDs). Covariates were considered adequately balanced when the SMDs were less than 0.1 for all covariates. Time-to-event analysis was conducted using Cox proportional hazards model. Baseline covariates were selected a priori based on clinical relevance and potential confounding. These included demographic characteristics, comorbidities, and clinical variables. Covariates used in the propensity score models are shown in Tables 1 and 2. We evaluated the dose-response effect of the β3 agonists by comparing 25 mg vs 50 mg drug regimens. Subgroup analyses were performed across key demographic and clinical strata.

**Table 1.**
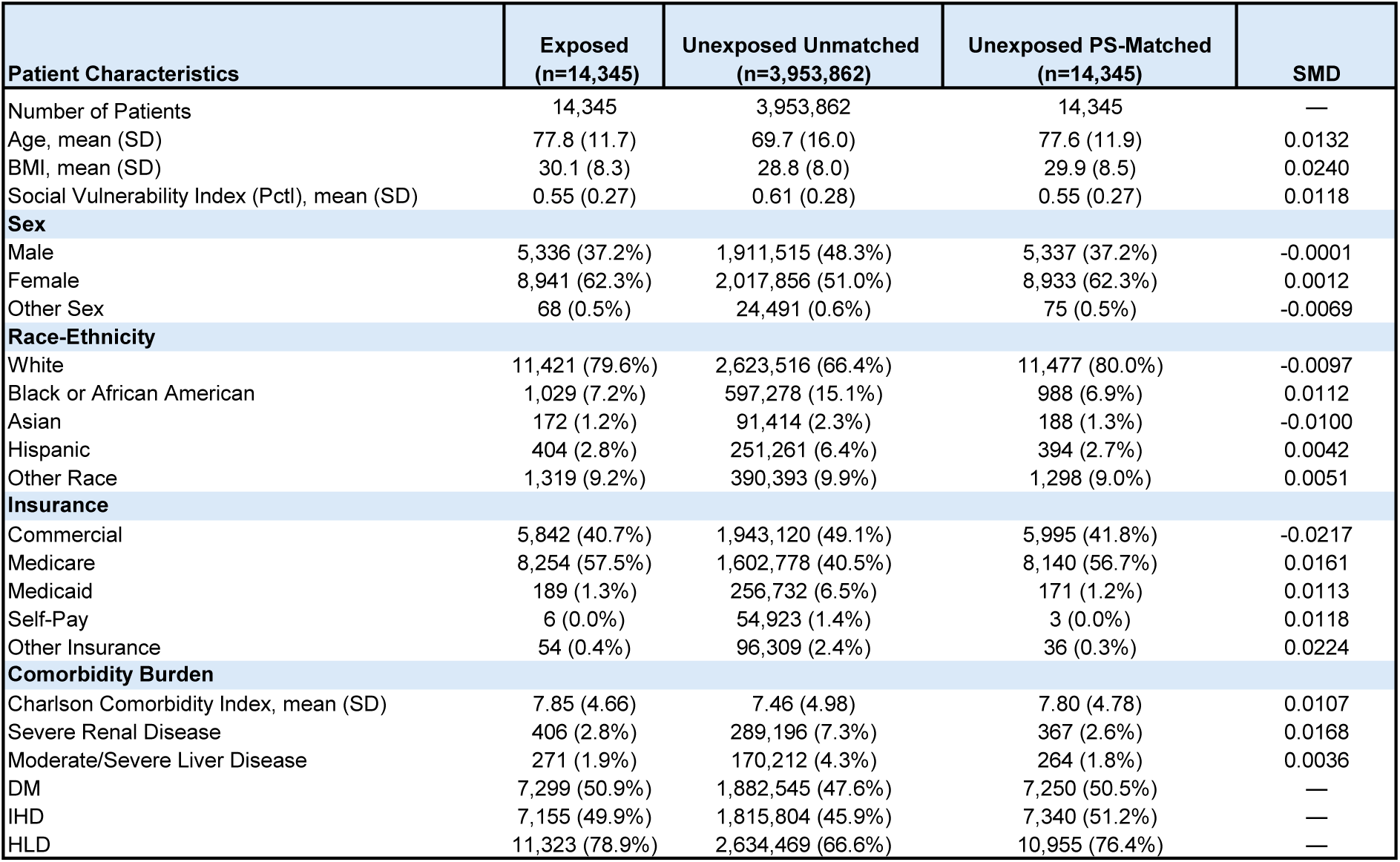
Demographics of control patients and those exposed to a β3-agonist before and after propensity score matching. BMI, body mass index; SVI, Social Vulnerability Index; DM, diabetes mellitus; IHD, ischemic heart disease; HLD, hyperlipidemia; SMD, standardized mean difference; PS, propensity score.

| Patient Characteristics | Exposed<br>(n=14,345) | Unexposed Unmatched<br>(n=3,953,862) | Unexposed PS-Matched<br>(n=14,345) | SMD |
| --- | --- | --- | --- | --- |
| Number of Patients | 14,345 | 3,953,862 | 14,345 | — |
| Age, mean (SD) | 77.8 (11.7) | 69.7 (16.0) | 77.6 (11.9) | 0.0132 |
| BMI, mean (SD) | 30.1 (8.3) | 28.8 (8.0) | 29.9 (8.5) | 0.0240 |
| Social Vulnerability Index (Pctl), mean (SD) | 0.55 (0.27) | 0.61 (0.28) | 0.55 (0.27) | 0.0118 |
| **Sex** |  |  |  |  |
| Male | 5,336 (37.2%) | 1,911,515 (48.3%) | 5,337 (37.2%) | -0.0001 |
| Female | 8,941 (62.3%) | 2,017,856 (51.0%) | 8,933 (62.3%) | 0.0012 |
| Other Sex | 68 (0.5%) | 24,491 (0.6%) | 75 (0.5%) | -0.0069 |
| **Race-Ethnicity** |  |  |  |  |
| White | 11,421 (79.6%) | 2,623,516 (66.4%) | 11,477 (80.0%) | -0.0097 |
| Black or African American | 1,029 (7.2%) | 597,278 (15.1%) | 988 (6.9%) | 0.0112 |
| Asian | 172 (1.2%) | 91,414 (2.3%) | 188 (1.3%) | -0.0100 |
| Hispanic | 404 (2.8%) | 251,261 (6.4%) | 394 (2.7%) | 0.0042 |
| Other Race | 1,319 (9.2%) | 390,393 (9.9%) | 1,298 (9.0%) | 0.0051 |
| **Insurance** |  |  |  |  |
| Commercial | 5,842 (40.7%) | 1,943,120 (49.1%) | 5,995 (41.8%) | -0.0217 |
| Medicare | 8,254 (57.5%) | 1,602,778 (40.5%) | 8,140 (56.7%) | 0.0161 |
| Medicaid | 189 (1.3%) | 256,732 (6.5%) | 171 (1.2%) | 0.0113 |
| Self-Pay | 6 (0.0%) | 54,923 (1.4%) | 3 (0.0%) | 0.0118 |
| Other Insurance | 54 (0.4%) | 96,309 (2.4%) | 36 (0.3%) | 0.0224 |
| **Comorbidity Burden** |  |  |  |  |
| Charlson Comorbidity Index, mean (SD) | 7.85 (4.66) | 7.46 (4.98) | 7.80 (4.78) | 0.0107 |
| Severe Renal Disease | 406 (2.8%) | 289,196 (7.3%) | 367 (2.6%) | 0.0168 |
| Moderate/Severe Liver Disease | 271 (1.9%) | 170,212 (4.3%) | 264 (1.8%) | 0.0036 |
| DM | 7,299 (50.9%) | 1,882,545 (47.6%) | 7,250 (50.5%) | — |
| IHD | 7,155 (49.9%) | 1,815,804 (45.9%) | 7,340 (51.2%) | — |
| HLD | 11,323 (78.9%) | 2,634,469 (66.6%) | 10,955 (76.4%) | — |

#### Bacterial Colony Forming Unit (CFU) Enumeration

The bacterial burden in the peritoneum, blood, liver, and spleen following CLP was quantified using standard plating technique per the American Society for Microbiology (57). Briefly, blood, peritoneal lavage, liver, and spleen were collected from mice in sterile Eppendorf tubes and kept in ice. Whole spleens and a piece of liver were harvested and weighed prior to being placed in bead homogenizing tubes (Omni International, Kennesaw, Georgia). For these organs, 1 mL of sterile phosphate-buffered saline (PBS) was added, and samples were homogenized using a BeadBug 6 MicroTube Homogenizer (Research Products International, Mount Prospect, IL). Homogenization consisted of two cycles at 4,350 RPM for 30 seconds each, separated by a 30 second rest period. All samples were stored on ice to arrest further bacterial growth. All samples (blood, peritoneal lavage fluid, and organ lysate) were plated on tryptic soy agar plates with 5% sheep’s blood (Thermo Scientific, Waltham, MA) and incubated under static conditions at 37 °C with 5% CO_2_ for 18 h to allow aerobic bacterial growth using standard dilution methods (57). Following incubation, samples were counted for CFU enumeration. Final counts normalized to organ mass prior to processing and expressed as CFU/g. Based on volumes of tissue and fluids obtained at necropsy, the limits of detection for blood and peritoneal lavage fluid were 1.7 log_10_ CFU/mL, 1.6 log_10_ CFU/g for liver, and 2.5 log_10_ CFU/g for spleen.

#### Flow Cytometry data

Single-cell suspensions were prepared from peritoneal lavage fluid, spleens, and mesenteric lymph nodes (MLNs). For peritoneal cells, the peritoneal cavity was lavaged with 5 mL PBS, gently massaged, and the fluid was collected. Spleens and MLNs were mechanically dissociated through a 100 μm cell strainer. Splenocytes were incubated with RBC lysis buffer (Thermo Fisher Scientific) at room temperature for 5 minutes and washed with PBS. Cells were resuspended in blocking buffer containing 0.5% BSA and anti-CD16/CD32 antibody (Thermo Fisher Scientific, 14-0161-81). Then, cells were stained with fluorophore-conjugated antibodies (see table below) and a viability dye (Zombie R718 fixable viability dye, BioLegend, 423115) at 4°C for 20 minutes in the dark, followed by fixation with BD Cytofix/Cytoperm (BD Biosciences, #554714) at 4°C for 20 minutes in the dark. Flow cytometry data were acquired on an Aurora spectral flow cytometer (Cytek Biosciences) using SpectroFlo software (version 3.3.0). Data analysis was performed using FlowJo (version 11).

### Antibodies used for Flow Cytometry

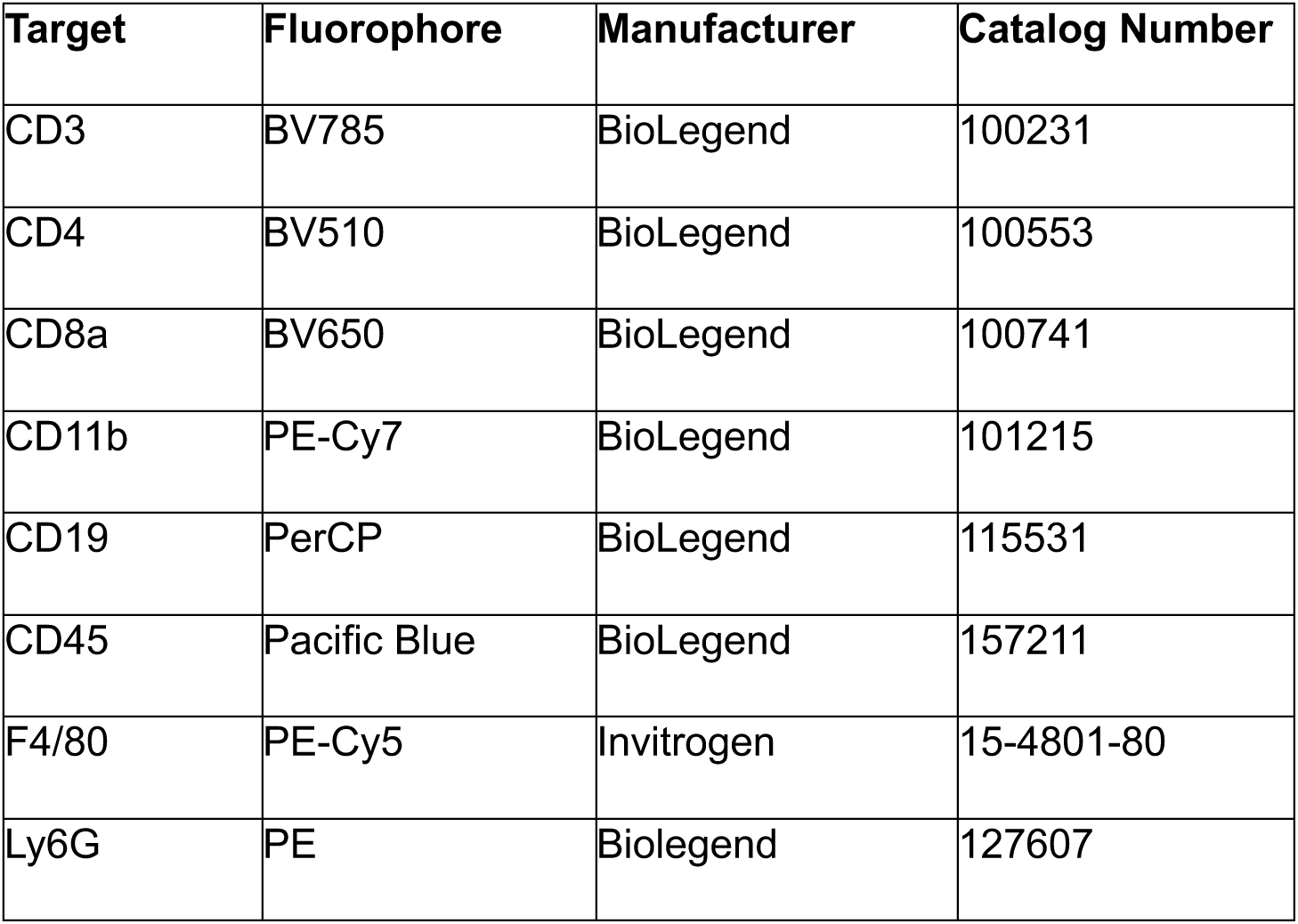

#### Metabolic Studies

Blood glucose, ketone, non-esterified fatty acid (NEFA) and glycerol measurements were performed on on whole blood obtained by tail vein prick. Glucose and ketones were measured using the Keto-Mojo GK+ meter, and NEFAs and glycerol were measured using LabAssay NEFA kit-Fujifilm and Free Glycerol Reagent (Sigma) respectively.

#### Serum Cytokines

Cytokine concentrations in mouse serum were measured using a bead-based multiplex immunoassay performed in a 96-well plate format according to the manufacturer’s instructions (Bio-Rad). The plate was analyzed using a Bio-Plex 200 multiplex reader (Bio-Rad Laboratories, Hercules, CA). Data acquisition and analysis were performed using Bio-Plex Manager Software version 6.2. Standard curves were generated by plotting mean fluorescence intensity (MFI) versus known concentrations of each cytokine, and sample concentrations (pg/mL) were determined by interpolation from these curves.

#### Immunoblot Aanalysis

Snap frozen tissues were homogenized in 1X RIPA buffer in a beadmill with addition of protease and phosphatase inhibitors (Thermo Fisher Scientific). After homogenization, total protein was estimated by using BCA Protein Assay Kit (Thermo Scientific). Lysates were then prepared in 1X SDS in RIPA lysis buffer and separated on Tris-HCl protein gel (Bio-Rad) and transferred to a polyvinylidene fluoride membranes (Thermo Fisher Scientific). They were then blocked in 5% blotting-grade milk, incubated in appropriate primary antibodies (1:1000 dilution) at 4°C overnight, and then in secondary HRP-conjugated antibody (Cytiva) for one hour at room temperature. Membranes were developed using chemiluminescent ECL reagents (Thermo Fisher Scientific) and images were collected by using ChemiDoc and Image Lab (Bio-Rad).

The following antibodies were used: anti-phospho-ERK1/2 (1:1,000; 9101), anti-ERK1/2 (1:1,000; 4695), anti-phospho-HSL (1:1,000, Ser660; 4126), anti-phospho-PKA substrate (1:1,000; 9624), anti-HSL (1:1,000; 4107), anti-p38 MAPK (1:1,000;9212), anti-phosphop-p38 (1:1,000;4511), anti-phospho-STAT3 Tyr705 (1:1000;9145) anti-STAT3 (1:1000;9139), anti-ATGL(1:1,000;2439) (all antibodies from Cell Signaling Technologies). Anti-HSP90 (1:1,000; 13171-1-AP), anti-αTubulin (1:1,000; 66031-1-IG) are from Proteintech.

#### Real-Time PCR Analysis

Snap frozen tissues were homogenized in Trizol using the bead-mill. Total RNA was isolated using E.Z.N.A. Total RNA Kit II (Omega Bio-Tek).1 μg RNA was reverse transcribed using the High-Capacity cDNA Reverse Transcription Kit (Thermo Fisher Scientific). Using 1 ng cDNA in a commercial SYBR Green PCR Master Mix (Thermo Fisher Scientific) and specific gene primers (table), qRT-PCR was performed using the QuantStudio 7 Pro Real-Time PCR System (Thermo Fisher Scientific) and normalized to the housekeeping gene Rn18s (referred to as *18s*). Analysis of qPCR data was conducted via the 2-ΔΔCT method.

#### Primers used for qRT-PCR

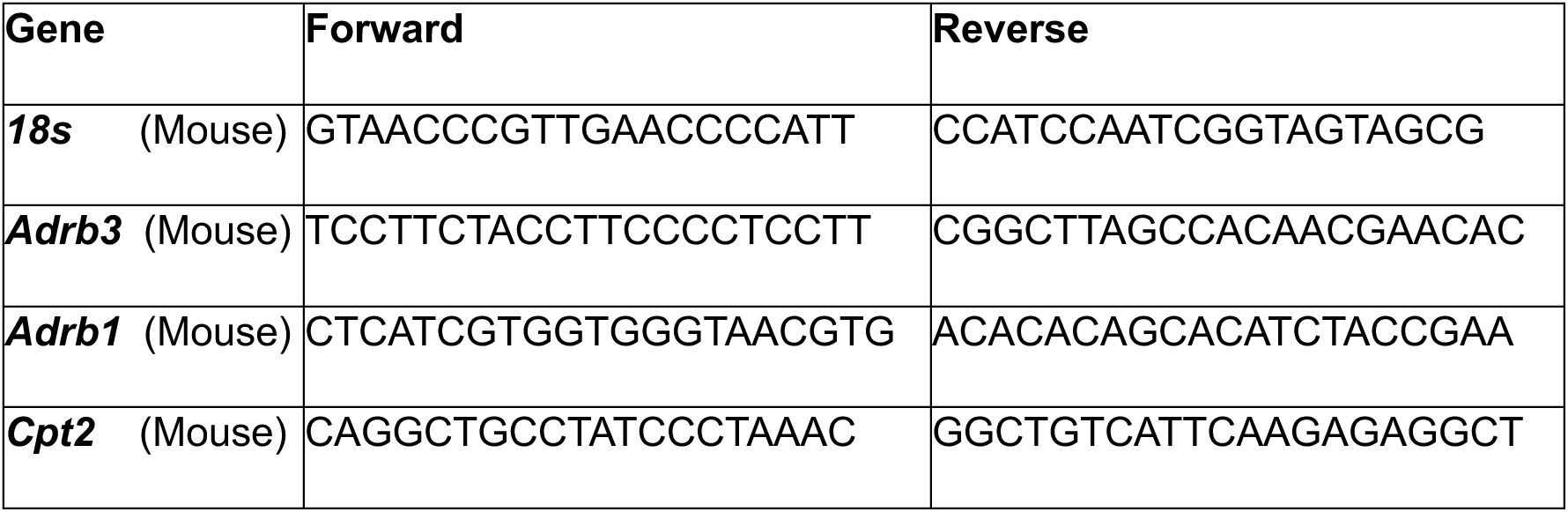

---

**Figure S1.**
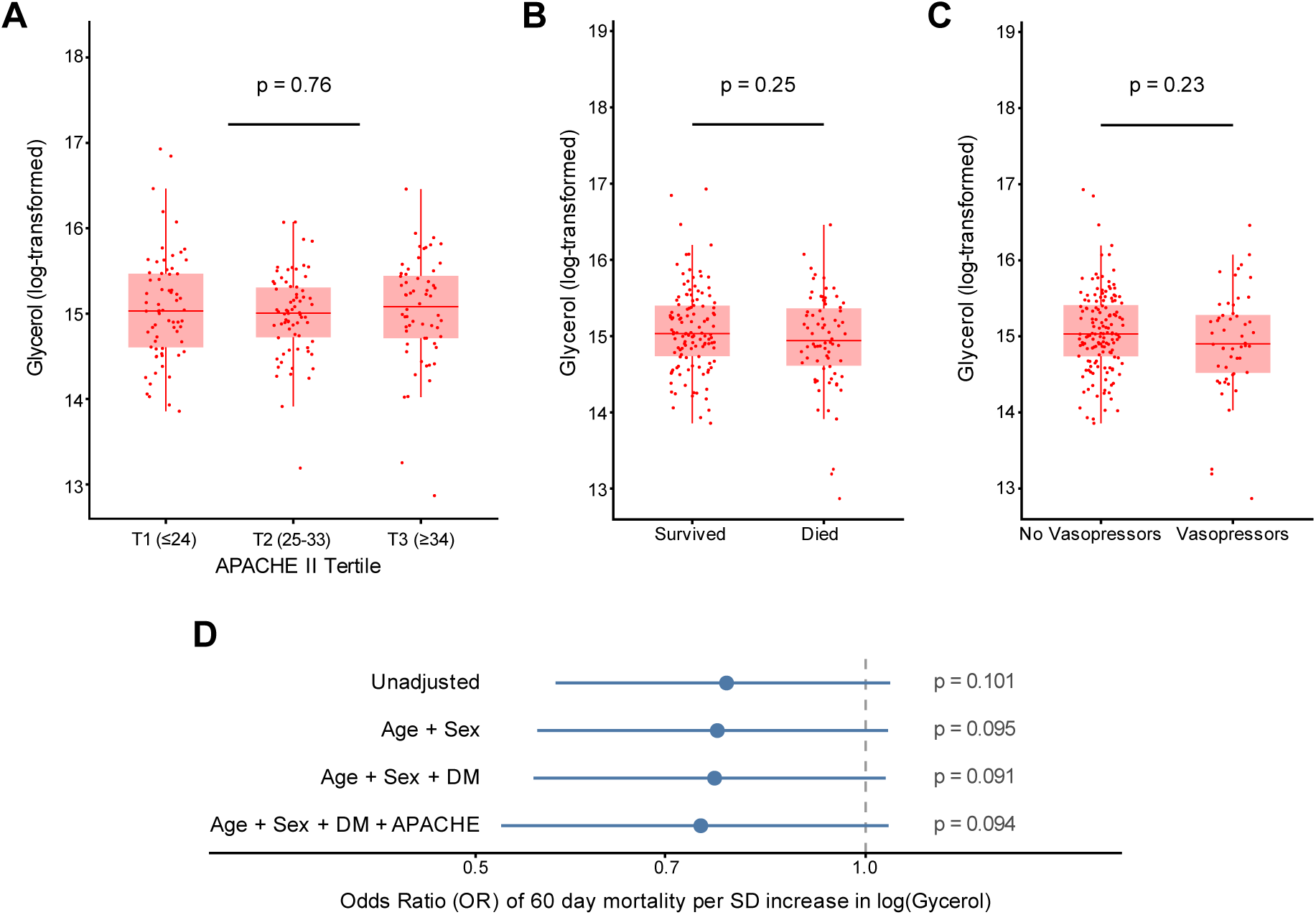
Circulating glycerol does not correlate with disease severity and survival in human sepsis. **(A)** Glycerol score stratified by Acute Physiology and Chronic Health Evaluation (APACHE II) tertiles. **(B)** Glycerol score stratified by 60-day mortality status. **(C)** Glycerol score stratified by need for vasopressors at enrollment. **(D)** Forest plot of odds ratios (ORs) for 60-day mortality from logistic regression models for glycerol score, per standard deviation increase, adjusted for age, sex, and diabetes. In panels **A–C**, horizontal bars indicate group means with error bars indicating standard error of the mean (SEM); significance was assessed by Wilcoxon rank-sum test (two-group comparisons) or pairwise comparison between T1 and T3; *p < 0.05, **p < 0.01, ***p < 0.001, ****p < 0.0001.

**Figure S2:**
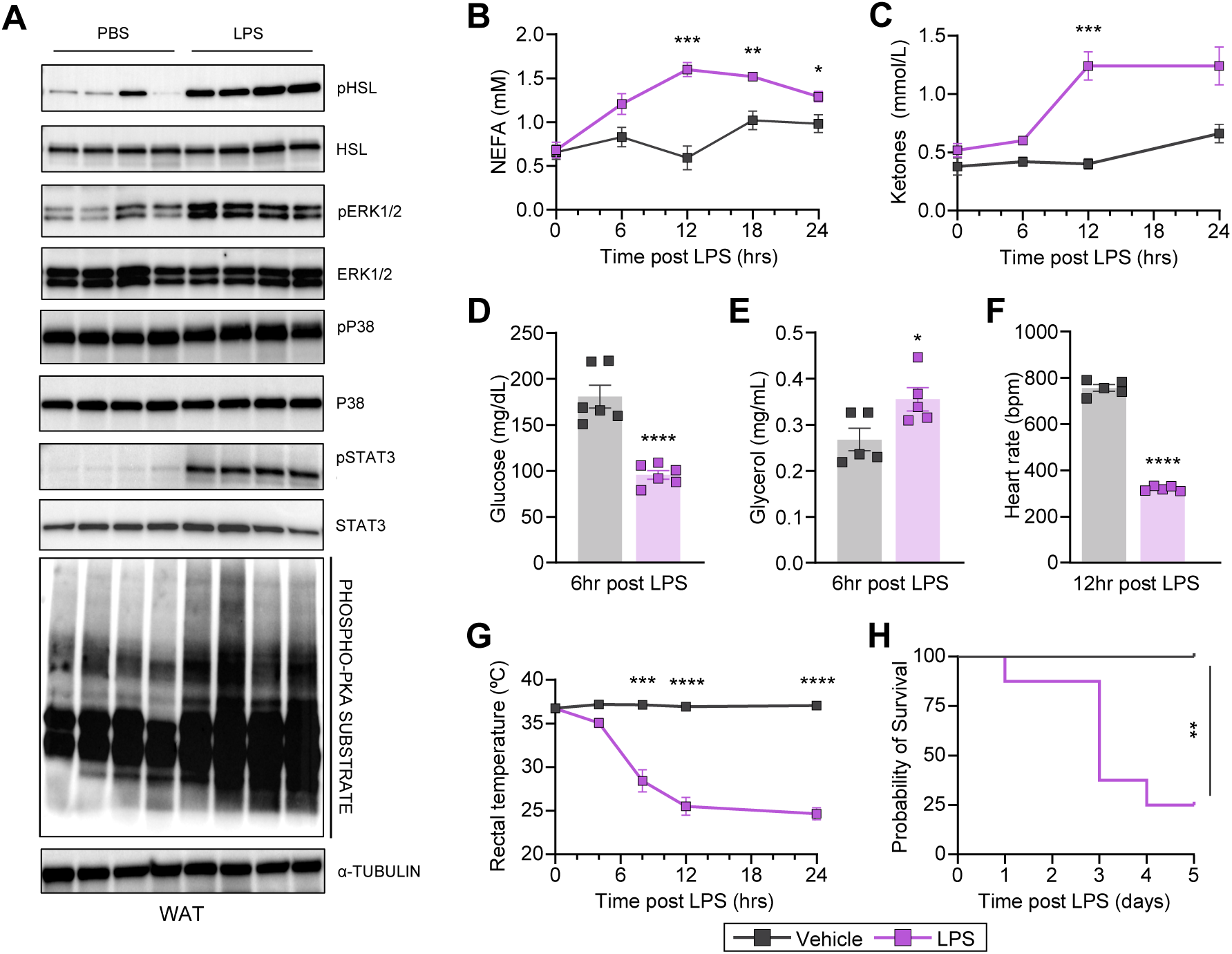
**Inflammation triggers adipose lipolysis and a conserved metabolic stress response**. **(A)** Immunoblot of pHSL, HSL, pERK, ERK, pSTAT3, STAT3, pP38, P38 and phospho-PKA substrate from WAT after LPS (n=4/group). Circulating **(B)** non-esterified fatty acids (NEFAs) (n=5/group), **(C)** ketones (n=5/group), **(D)** glucose (n=6/group), and **(E)** glycerol level (n=5/group); and **(F)** heart rates (n=5/group), **(G)** rectal temperatures (n=5/group), and **(H)** Kaplan-Meier survival curve (n=8/group) after lipopolysaccharide (LPS). LPS dose was 10 mg/kg Data are represented as mean ± SEM. *p < 0.05; **p<0.01; ***p < 0.001; ****p < 0.0001, log-rank (Mantel–Cox) test.

**Figure S3:**
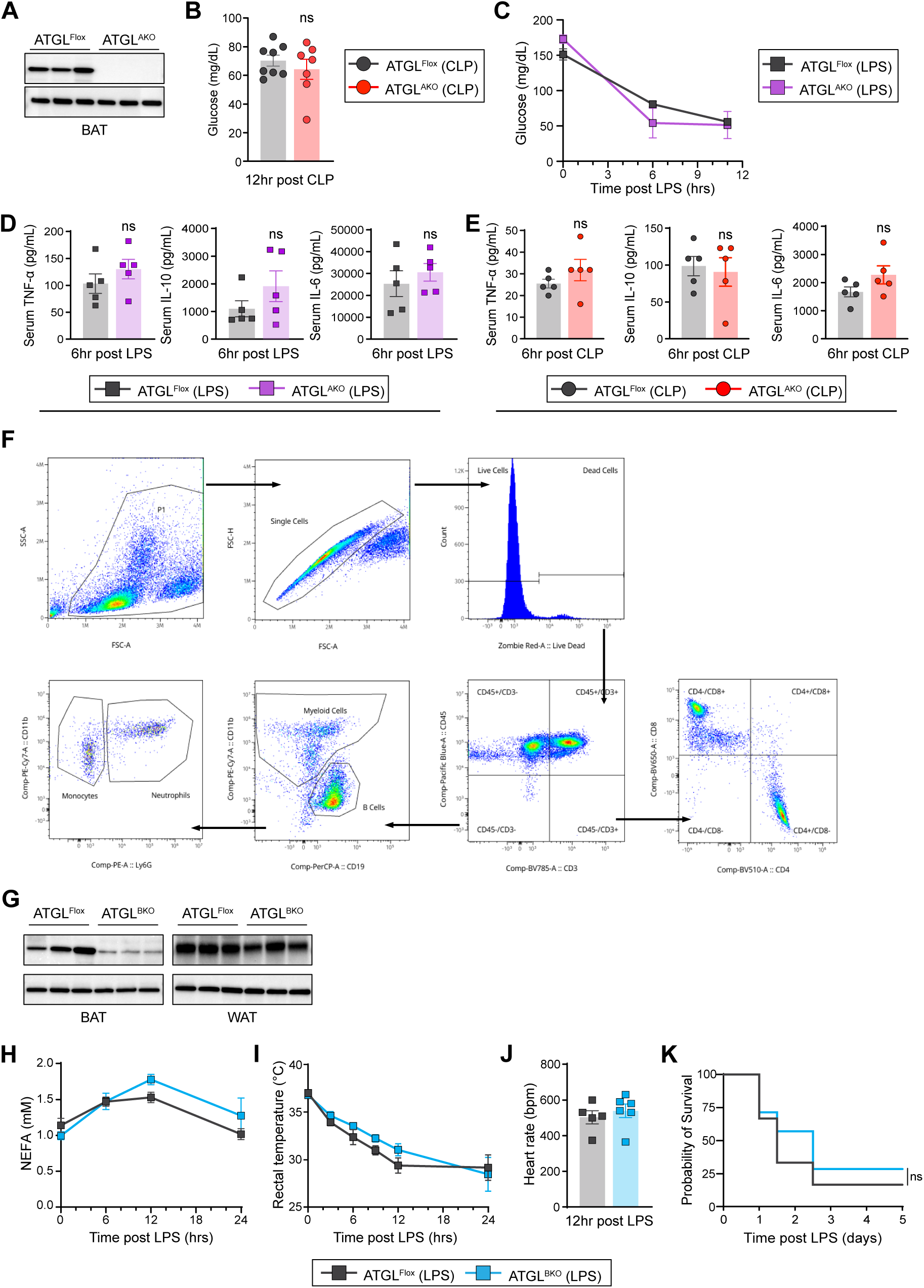
White adipose tissue lipolysis is necessary for tolerance to infection. **(A)** Immunoblot of ATGL expression in BAT (n=3/group). Circulating glucose (n=7/group) after **(B)** cecal ligation and puncture (CLP) and **(C)** lipopolysaccharide (LPS). Serum TNF-α, IL-10, and IL-6 after **(D)** LPS (n=5/group) and **(E)** CLP (n=5/group). **(F)** Gating strategy for flow cytometry analysis (Myeloid cells: CD11b+CD45+, B cells: CD19+CD45+, Neutrophils (PMN): CD11b+Ly6G+CD45+, Monocytes: CD11b+Ly6G−CD45+, T cells: CD3+CD45+, CD4+ T cells: CD3+CD4+CD45+, and CD8+ T cells: CD3+CD8+CD45+). **(G)** Immunoblot of ATGL in BAT and WAT (n=3/group). **(H)** Circulating NEFAs, **(I)** rectal temperatures, **(J)** heart rates, and **(K)** Kaplan–Meier survival curve after LPS. For (**B** and **H–K**), the LPS dose was 5 mg/kg. Data are represented as mean ± SEM. *p < 0.05; **p<0.01; ***p < 0.001; ****p < 0.0001, log-rank (Mantel–Cox) test.

**Figure S4:**
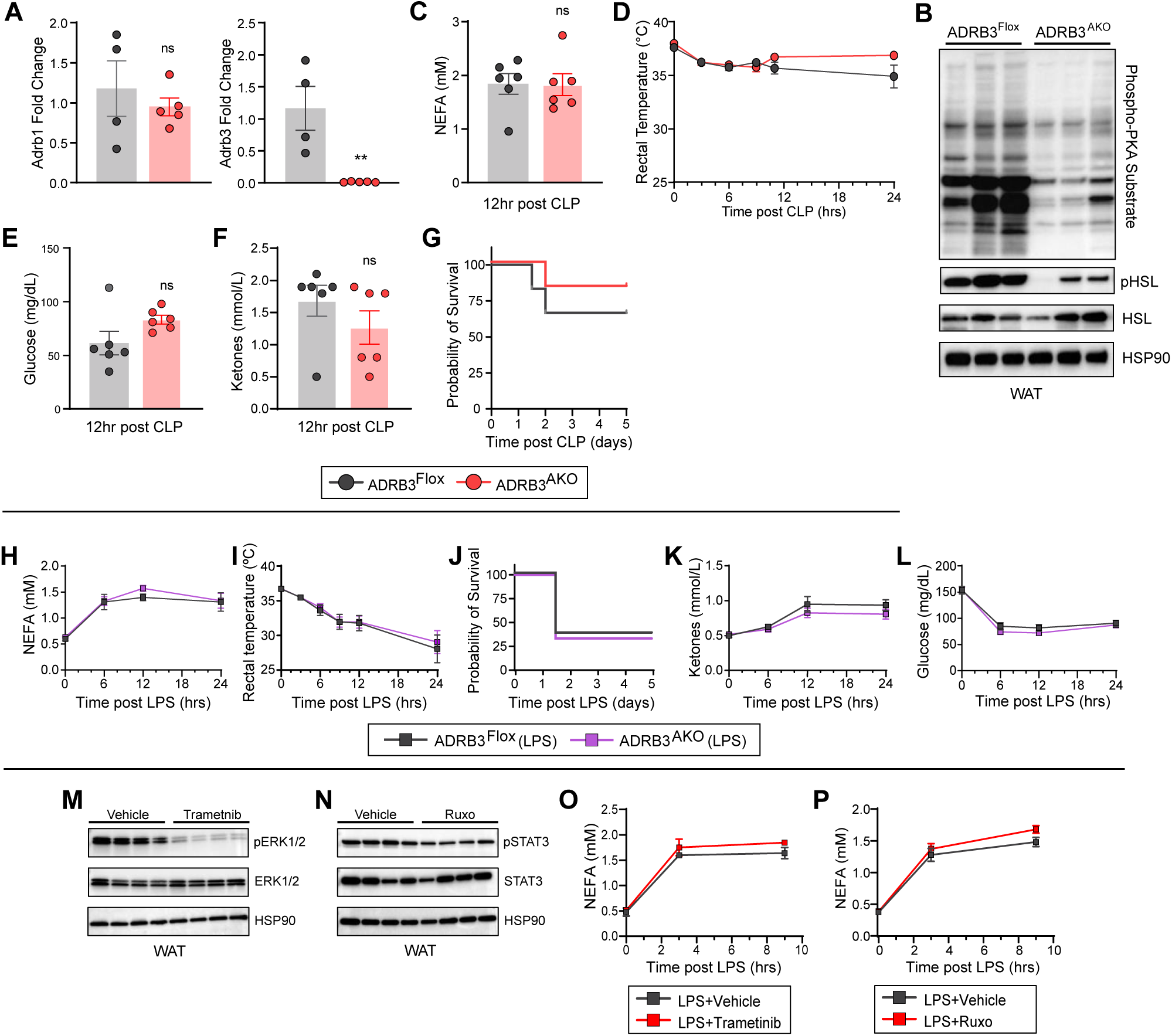
Infection-induced adipose lipolysis is redundantly regulated. **(A)** Expression of *Adrb1* and *Adrb3* from white adipose tissue (WAT) (n=4/group). **(B)** Immunoblot of phospho-PKA substrate, pHSL and HSL from WAT 3 h after liposaccharide (LPS) (n=3/group). **(C)** Circulating non-esterified fatty acids (NEFAs), after **(D)** rectal temperatures after CLP. Circulating **(E)** glucose and **(F)** ketones and **(G)** Kaplan–Meier survival curve after cecal ligation and puncture (CLP) (**C-G**, n=6/group). Circulating **(H)** NEFAs, **(I)** rectal temperatures, **(J)** Kaplan–Meier survival curve, **(K)** ketones and **(L)** glucose after LPS (**H-L**, LPS dose was 7.5 mg/kg and n=6-8/group). **(M)** Immunoblot of pERK1/2 and ERK1/2 in WAT (n=4/group) and **(N)** Immunoblot of pSTAT3 and STAT3 in the WAT (n=4/group). **(O)** and **(P)** circulating post LPS (2.5 mg/kg) NEFAs after Trametinib and Ruxolitinib respectively. Data are represented as mean ± SEM. *p < 0.05; **p<0.01; ***p < 0.001; ****p < 0.0001, log-rank (Mantel–Cox) test.

**Figure S5:**
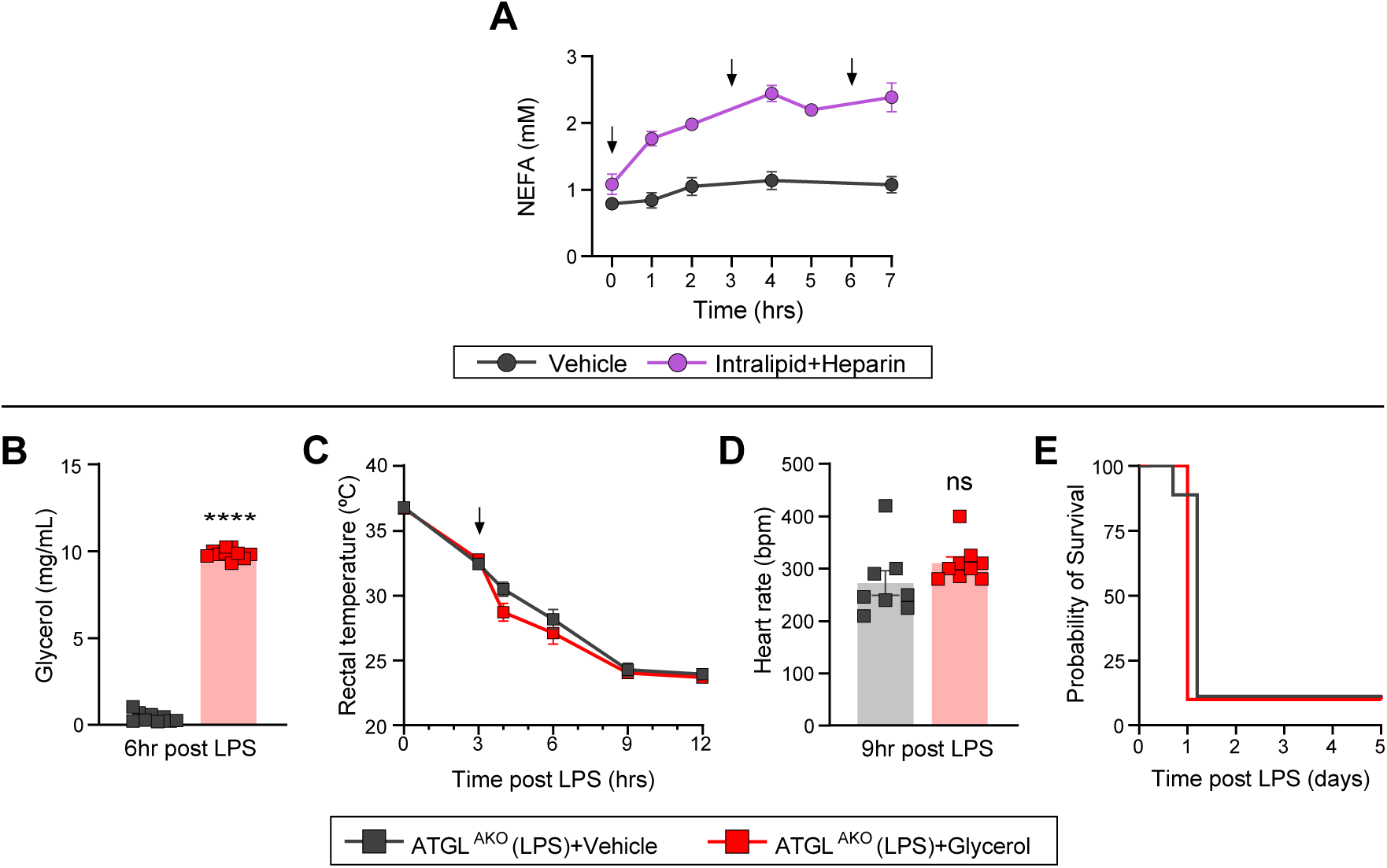
Circulating glycerol does not mediate tolerance to infection. **(A)** Circulating non-esterified fatty acids (NEFAs) after intralipid heparin supplementation. **(B)** Circulating glycerol, **(C)** rectal temperatures, **(D)** heart rates, and **(E)** Kaplan–Meier survival curve after LPS with glycerol supplementation. For (B-E) LPS dose was 5 mg/kg and n=9-10/group. Data are represented as mean ± SEM. *p < 0.05; **p<0.01; ***p < 0.001; ****p < 0.0001, log-rank (Mantel–Cox) test.

**Figure S6:**
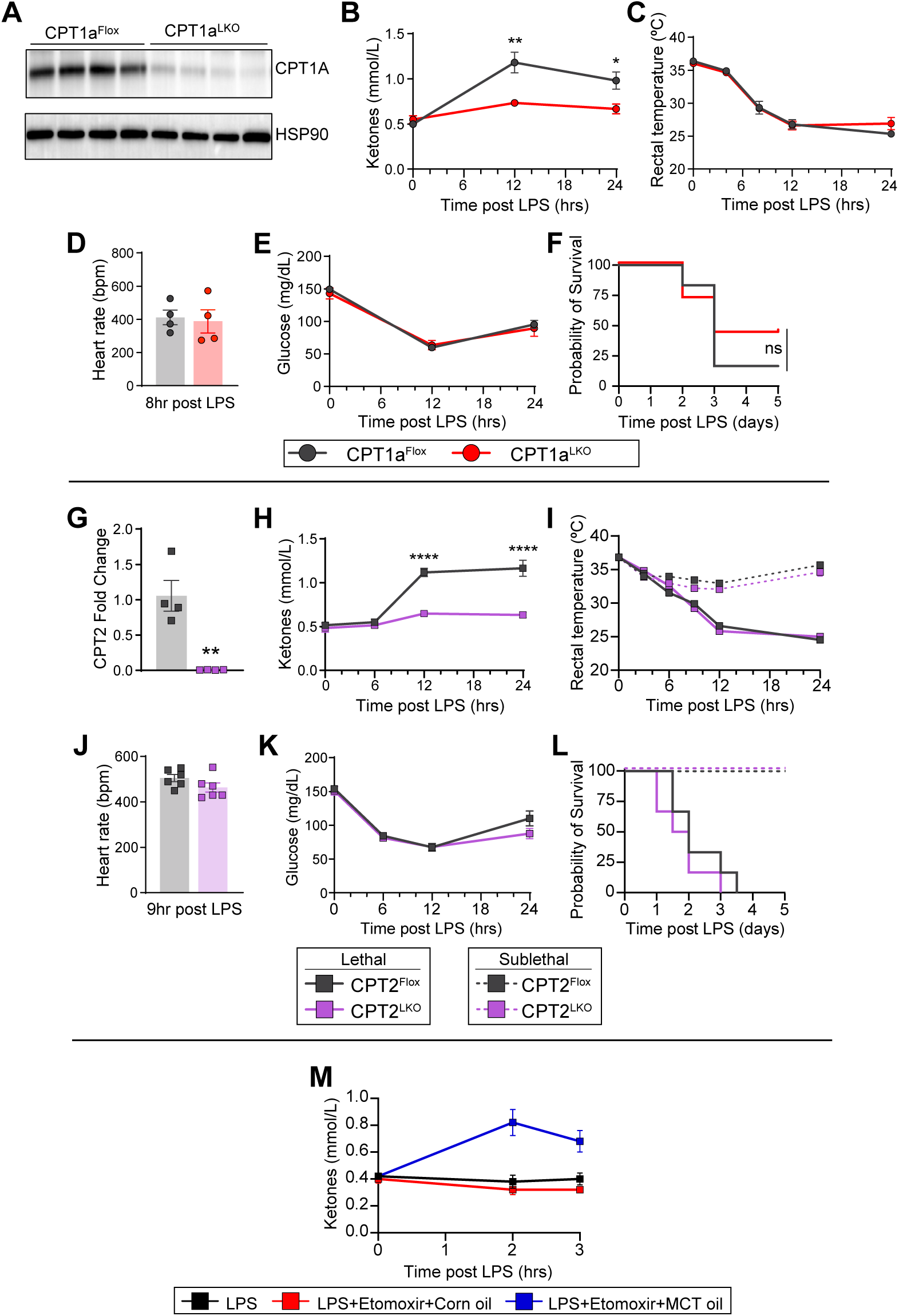
Hepatic fatty acid oxidation is not required for tolerance to inflammation. **(A)** Immunoblot of CPT1A in liver (n=4/group). Circulating **(B)** ketones (n=6-7/group), **(C)** rectal temperatures (n=6-7/group); and **(D)** heart rates (n=4/group), **(E)** glucose (n=6-7/group), and **(F)** Kaplan–Meier survival curve (n=6-7/group) after lipopolysaccharide (LPS) (5 mg/kg). **(G)** Expression of *Cpt2* in liver (n=4). Circulating **(H)** ketones, **(I)** rectal temperatures, **(J)** heart rates, **(K)** glucose, and **(L)** Kaplan–Meier survival curve after LPS. **(M)** Circulating ketones after LPS (5 mg/kg) and etomoxir with medium-chain triglyceride (MCT) or corn oil supplementation (n=5/group). For (**I** and **L**), LPS dose was 5 mg/kg (lethal) and 2.5 mg/kg (Sublethal) (n=6/group). For H, J, and K LPS dose was 5 mg/kg. Data are represented as mean ± SEM. *p < 0.05; **p < 0.01; ***p < 0.001; ****p < 0.0001, log-rank (Mantel–Cox) test.

**Figure S7:**
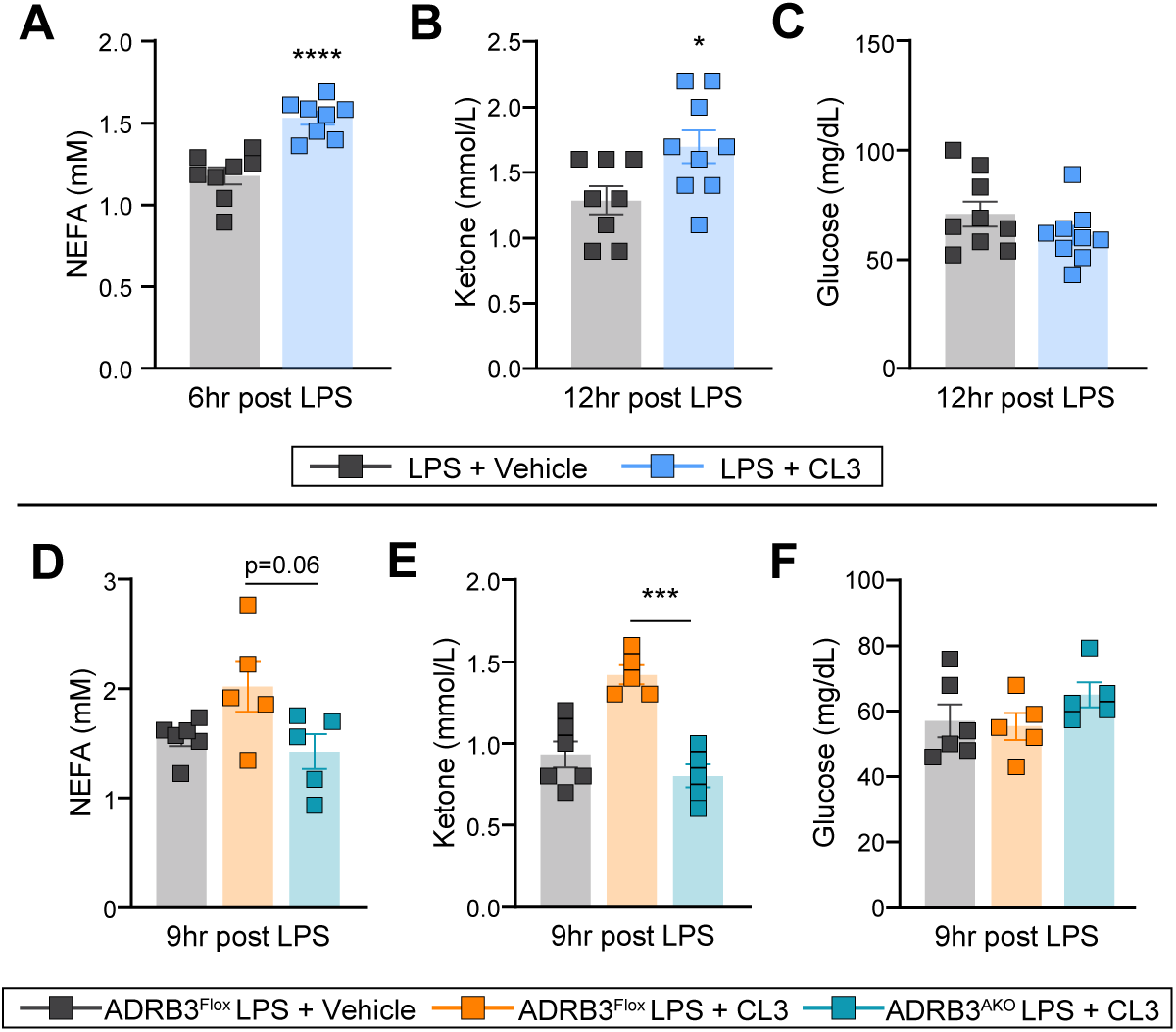
CL316,243 promotes tolerance through the adipose β3 receptor. Circulating. **(A)** non-esterified fatty acids (NEFAs), **(B)** ketones, and **(C)** glucose levels after lipopolysaccharide (LPS) and CL316,243 (CL3). For (A-C) n=8/group. Circulating **(D)** NEFAs, **(E)** ketones, and **(F)** glucose levels after LPS and CL3. For (D-F), n=5-6/group. LPS dose was 10 mg/kg. Data are represented as mean ± SEM. *p < 0.05; **p < 0.01; ***p < 0.001; ****p < 0.0001.

**Figure S8:**
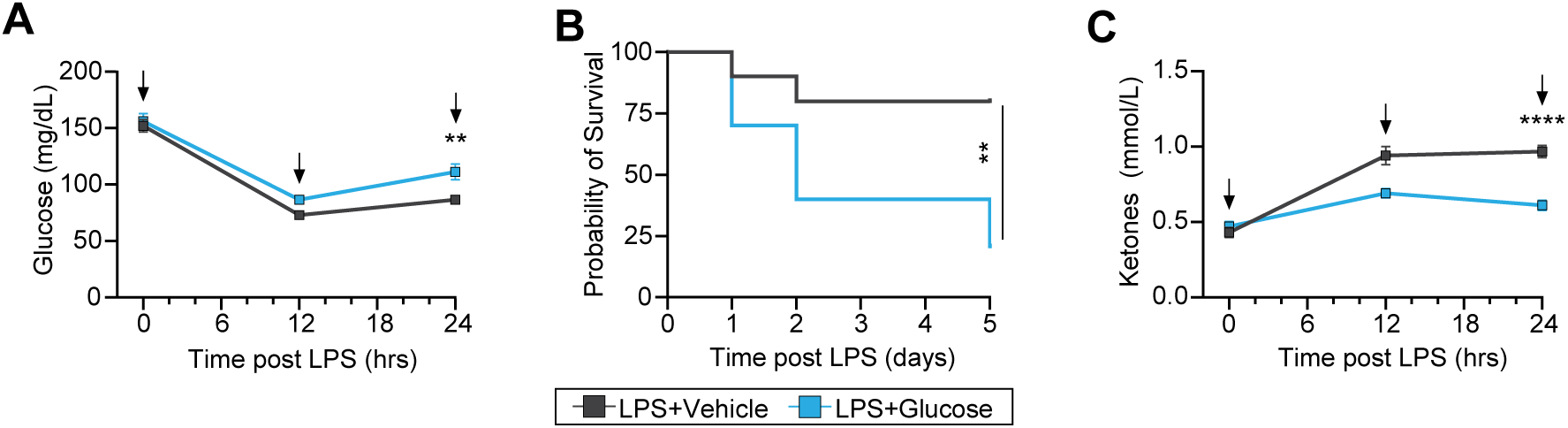
Glucose dissolved in water increases mortality with LPS. **(A)** Glucose, **(B)** Kaplan–Meier survival curve and **(C)** ketones after lipopolysaccharide (LPS) with glucose supplementation. Data are represented as mean ± SEM. *p < 0.05; **p < 0.01; ***p < 0.001; ****p < 0.0001, log-rank (Mantel–Cox) test. LPS dose was 10 mg/kg and n=10/group. Arrows indicate the time of CL3 administration and intralipid supplementation. Veh, vehicle. Vehicle was PBS, and glucose was dissolved in water.

**Table S1.**
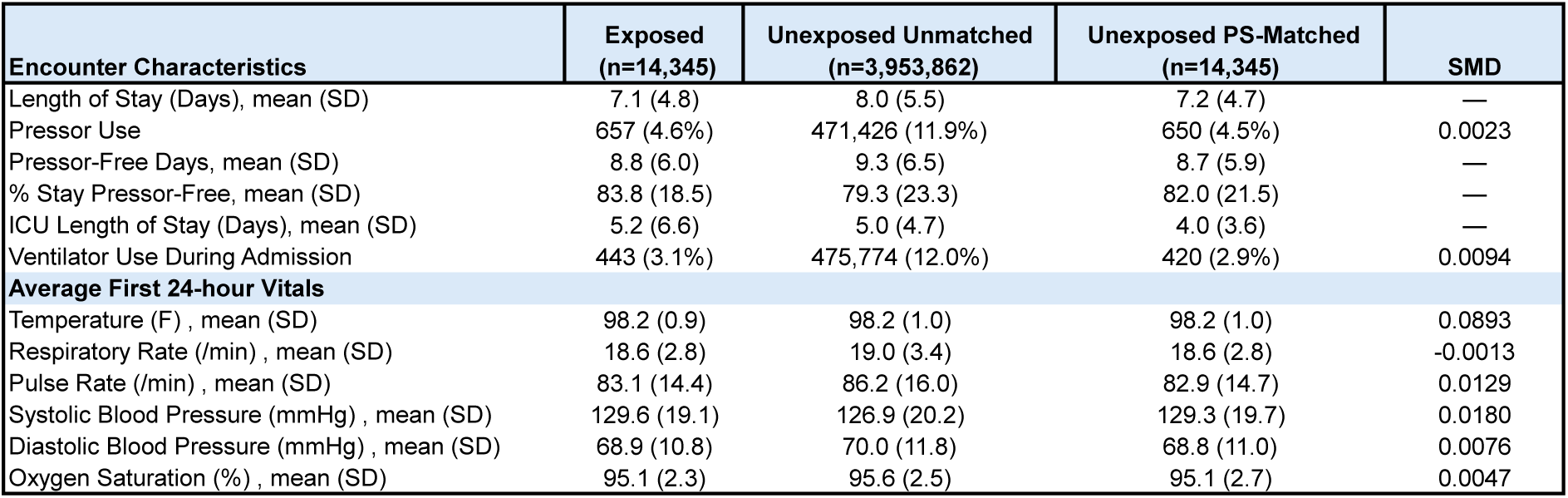
Encounter characteristics of control patients and those exposed to a β3-agonist before and after propensity score matching. ICU, intensive care unit; SMD, standardized mean difference; PS, propensity score.

