## Supplemental Data for "Lipid mobilization establishes metabolic tolerance and prevents autonomic collapse in infection"

**Antibodies used for Flow Cytometry**

| Target | Fluorophore | Manufacturer | Catalog Number |
|---|---|---|---|
| CD3 | BV785 | BioLegend | 100231 |
| CD4 | BV510 | BioLegend | 100553 |
| CD8a | BV650 | BioLegend | 100741 |
| CD11b | PE-Cy7 | BioLegend | 101215 |
| CD19 | PerCP | BioLegend | 115531 |
| CD45 | Pacific Blue | BioLegend | 157211 |
| F4/80 | PE-Cy5 | Invitrogen | 15-4801-80 |
| Ly6G | PE | Biolegend | 127607 |

**Primers used for qRT-PCR**

| Gene | Forward | Reverse |
| --- | --- | --- |
| ***18s*** (Mouse) | GTAACCCGTTGAACCCCATT | CCATCCAATCGGTAGTAGCG |
| ***Adrb3*** (Mouse) | TCCTTCTACCTTCCCCTCCTT | CGGCTTAGCCACAACGAACAC |
| ***Adrb1*** (Mouse) | CTCATCGTGGTGGGTAACGTG | ACACACAGCACATCTACCGAA |
| ***Cpt2*** (Mouse) | CAGGCTGCCTATCCCTAAAC | GGCTGTCATTCAAGAGAGGCT |

**Figure S1. Circulating glycerol does not correlate with disease severity and survival in human sepsis.**

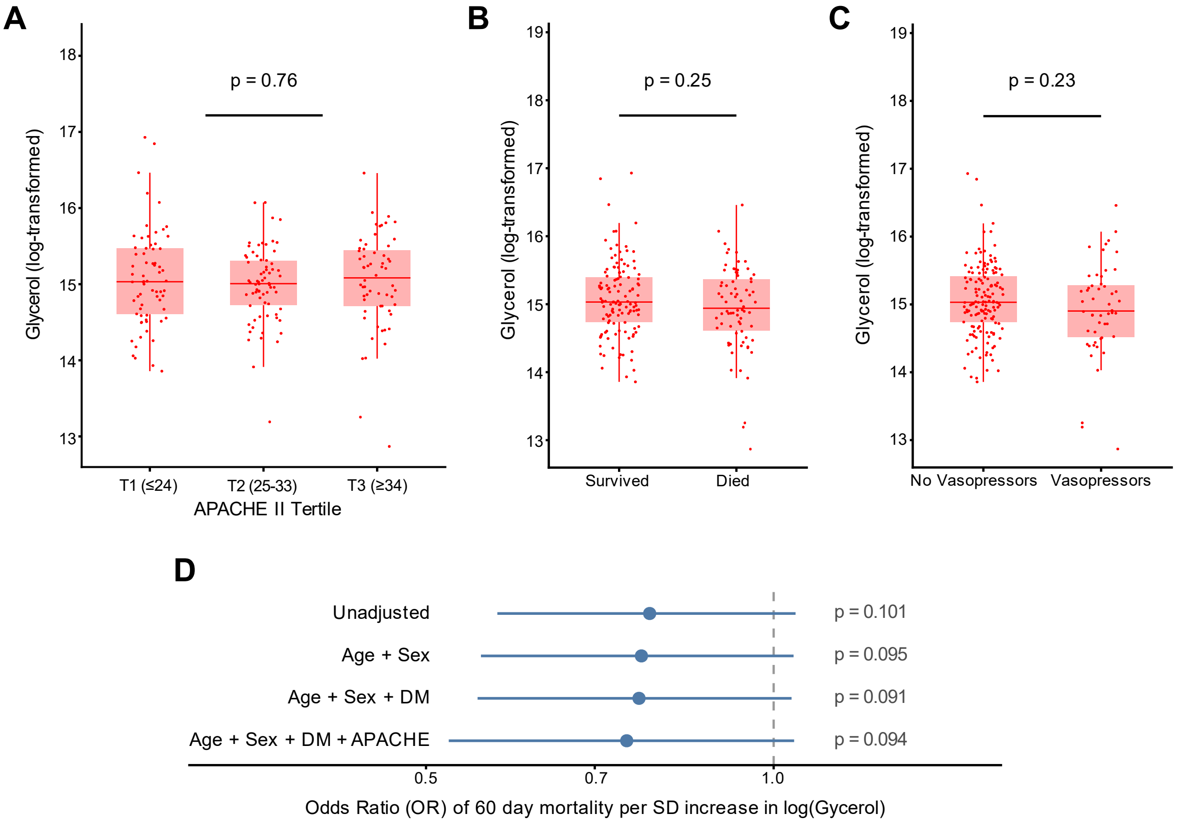

**Figure S1. Circulating glycerol does not correlate with disease severity and survival in human sepsis. (A)** Glycerol score stratified by Acute Physiology and Chronic Health Evaluation (APACHE II) tertiles. **(B)** Glycerol score stratified by 60-day mortality status. **(C)** Glycerol score stratified by need for vasopressors at enrollment. **(D)** Forest plot of odds ratios (ORs) for 60-day mortality from logistic regression models for glycerol score, per standard deviation increase, adjusted for age, sex, and diabetes. In panels **A–C**, horizontal bars indicate group means with error bars indicating standard error of the mean (SEM); significance was assessed by Wilcoxon rank-sum test (two-group comparisons) or pairwise comparison between T1 and T3; *$p < 0.05$, **$p < 0.01$, ***$p < 0.001$, ****$p < 0.0001$.

**Figure S2: Inflammation triggers adipose lipolysis and a conserved metabolic stress response.**

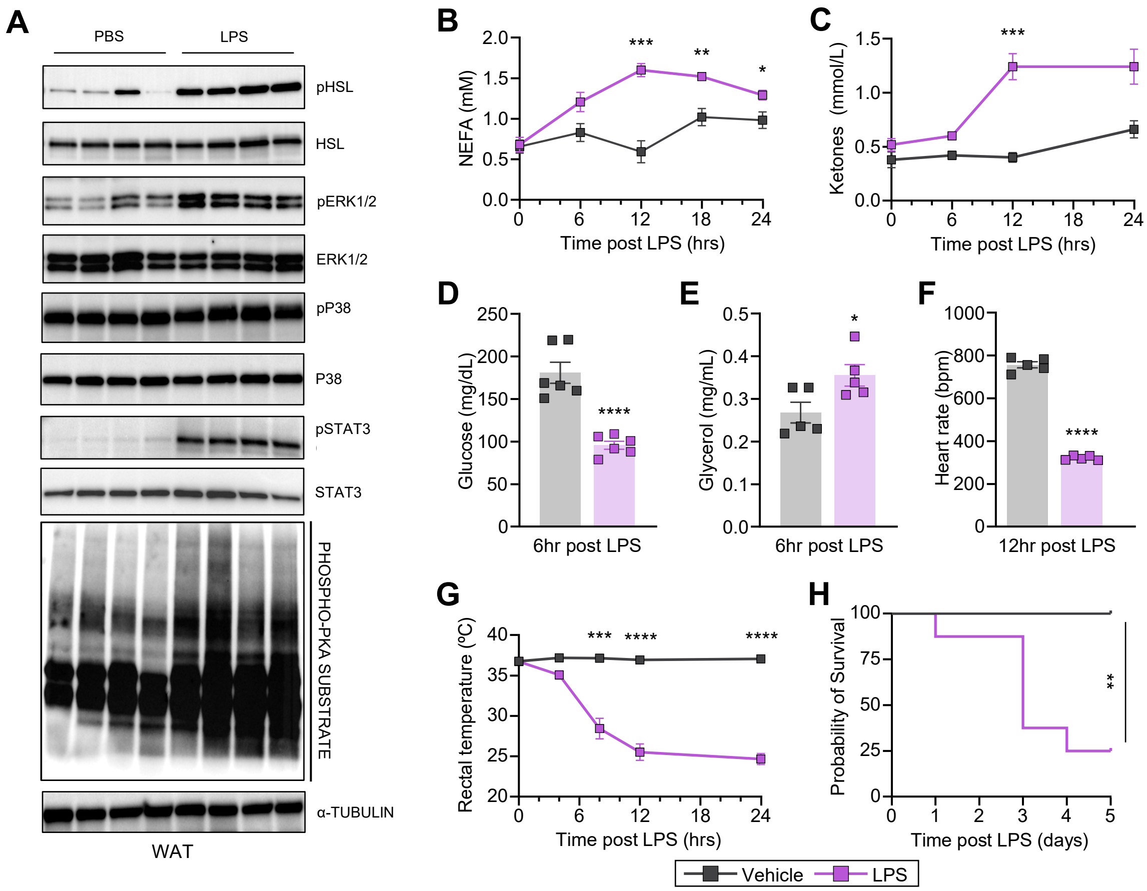

**Figure S2: Inflammation triggers adipose lipolysis and a conserved metabolic stress response**. **(A)** Immunoblot of pHSL, HSL, pERK, ERK, pSTAT3, STAT3, pP38, P38 and phospho-PKA substrate from WAT after LPS (n=4/group). Circulating **(B)** non-esterified fatty acids (NEFAs) (n=5/group), **(C)** ketones (n=5/group), **(D)** glucose (n=6/group), and **(E)** glycerol level (n=5/group); and **(F)** heart rates (n=5/group), **(G)** rectal temperatures (n=5/group), and **(H)** Kaplan-Meier survival curve (n=8/group) after lipopolysaccharide (LPS). LPS dose was 10 mg/kg Data are represented as mean ± SEM. *p < 0.05; **p<0.01; ***p < 0.001; ****p < 0.0001, log-rank (Mantel–Cox) test.

**Figure S3: White adipose tissue lipolysis is necessary for tolerance to infection.**

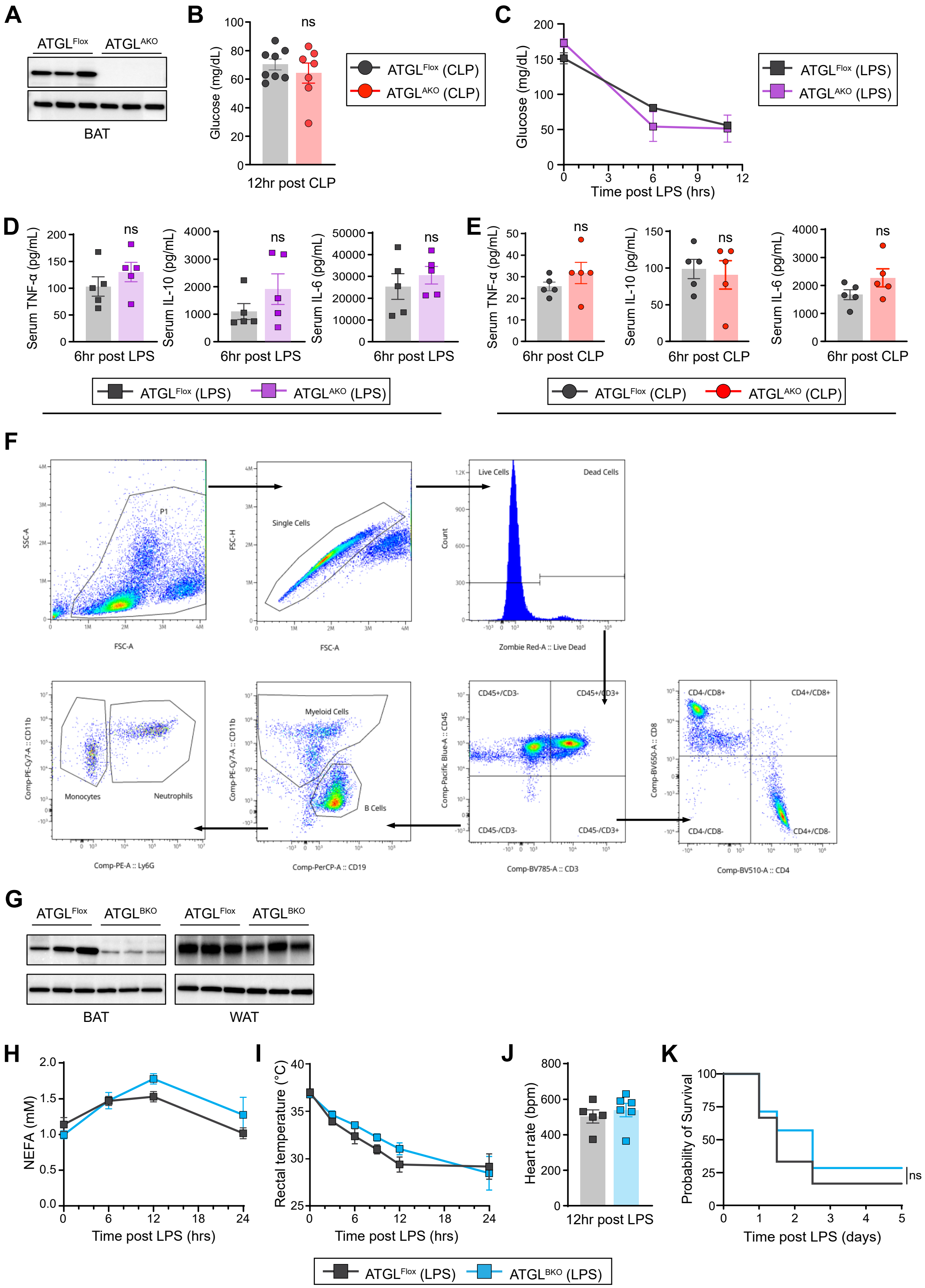

**Figure S3: White adipose tissue lipolysis is necessary for tolerance to infection. (A)** Immunoblot of ATGL expression in BAT (n=3/group). Circulating glucose (n=7/group) after **(B)** cecal ligation and puncture (CLP) and **(C)** lipopolysaccharide (LPS). Serum TNF-α, IL-10, and IL-6 after **(D)** LPS (n=5/group) and **(E)** CLP (n=5/group). **(F)** Gating strategy for flow cytometry analysis (Myeloid cells: CD11b+CD45+, B cells: CD19+CD45+, Neutrophils (PMN): CD11b+Ly6G+CD45+, Monocytes: CD11b+Ly6G−CD45+, T cells: CD3+CD45+, CD4+ T cells: CD3+CD4+CD45+, and CD8+ T cells: CD3+CD8+CD45+). **(G)** Immunoblot of ATGL in BAT and WAT (n=3/group). **(H)** Circulating NEFAs, **(I)** rectal temperatures, **(J)** heart rates, and **(K)** Kaplan–Meier survival curve after LPS. For (**B** and **H–K**), the LPS dose was 5 mg/kg. Data are represented as mean ± SEM. $*p < 0.05$; $**p<0.01$; $***p < 0.001$; $****p < 0.0001$, log-rank (Mantel–Cox) test.

**Figure S4: Infection-induced adipose lipolysis is redundantly regulated.**

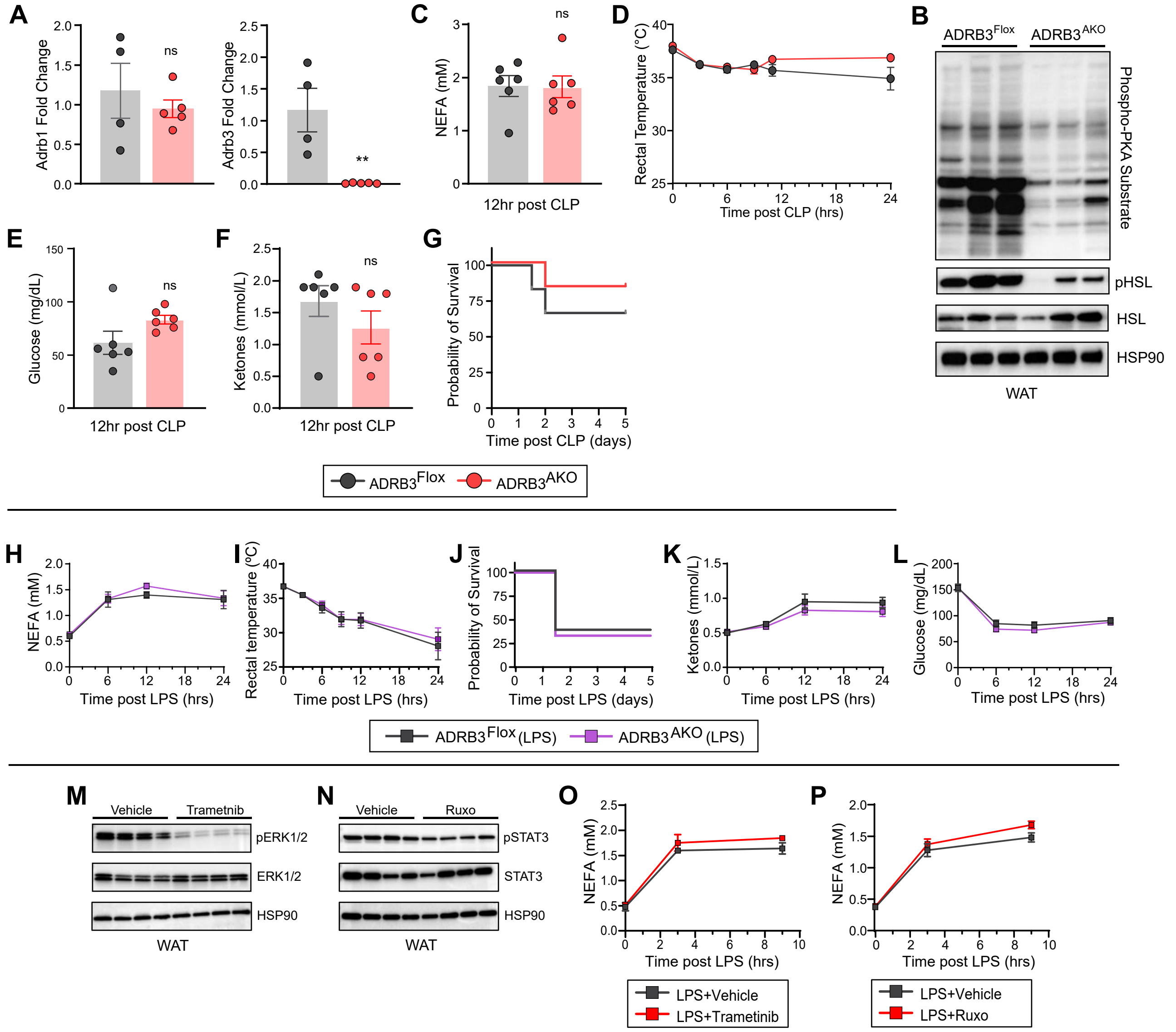

**Figure S4: Infection-induced adipose lipolysis is redundantly regulated. (A)** Expression of *Adrb1* and *Adrb3* from white adipose tissue (WAT) (n=4/group). **(B)** Immunoblot of phospho-PKA substrate, pHSL and HSL from WAT 3 h after liposaccharide (LPS) (n=3/group). **(C)** Circulating non-esterified fatty acids (NEFAs), after **(D)** rectal temperatures after CLP. Circulating **(E)** glucose and **(F)** ketones and **(G)** Kaplan–Meier survival curve after cecal ligation and puncture (CLP) (**C-G**, n=6/group). Circulating **(H)** NEFAs, **(I)** rectal temperatures, **(J)** Kaplan–Meier survival curve, **(K)** ketones and **(L)** glucose after LPS (**H-L**, LPS dose was 7.5 mg/kg and n=6-8/group). **(M)** Immunoblot of pERK1/2 and ERK1/2 in WAT (n=4/group) and **(N)** Immunoblot of pSTAT3 and STAT3 in the WAT (n=4/group). **(O)** and **(P)** circulating post LPS (2.5 mg/kg) NEFAs after Trametinib and Ruxolitinib respectively. Data are represented as mean ± SEM. *p < 0.05; **p<0.01; ***p < 0.001; ****p < 0.0001, log-rank (Mantel–Cox) test.

**Figure S5: Circulating glycerol does not mediate tolerance to infection.**

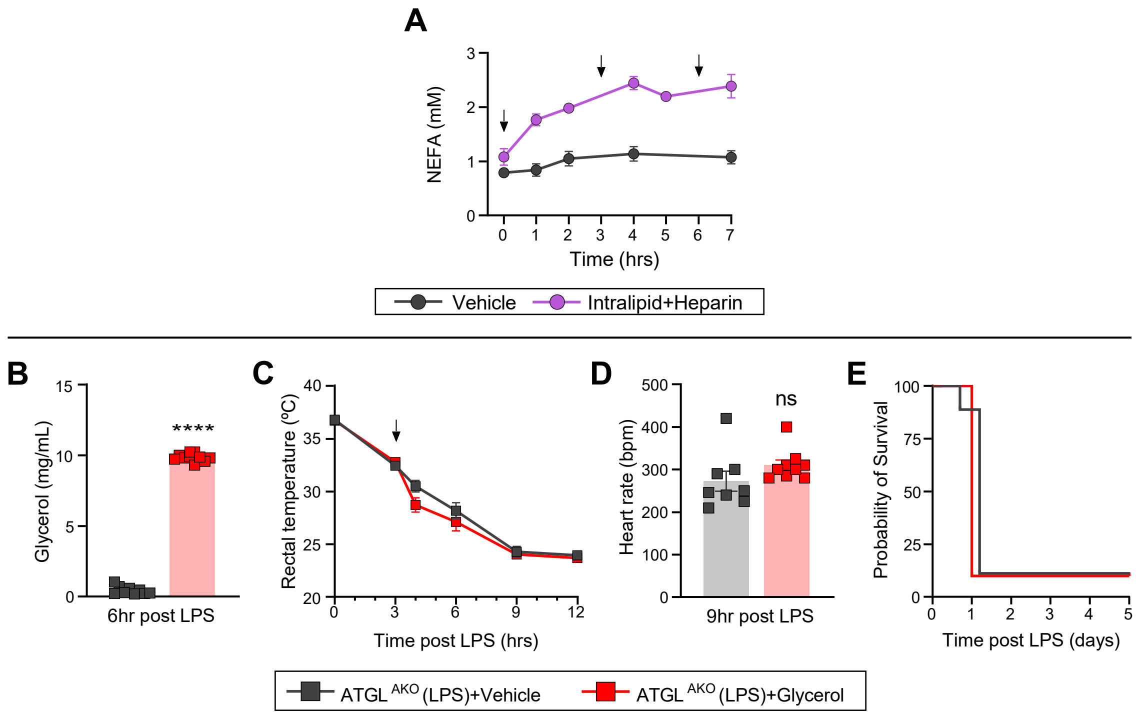

**Figure S5: Circulating glycerol does not mediate tolerance to infection. (A)** Circulating non-esterified fatty acids (NEFAs) after intralipid heparin supplementation. **(B)** Circulating glycerol, **(C)** rectal temperatures, **(D)** heart rates, and **(E)** Kaplan–Meier survival curve after LPS with glycerol supplementation. For (B-E) LPS dose was 5 mg/kg and n=9-10/group. Data are represented as mean ± SEM. *p < 0.05; **p<0.01; ***p < 0.001; ****p < 0.0001, log-rank (Mantel–Cox) test.

**Figure S6: Hepatic fatty acid oxidation is not required for tolerance to inflammation.**

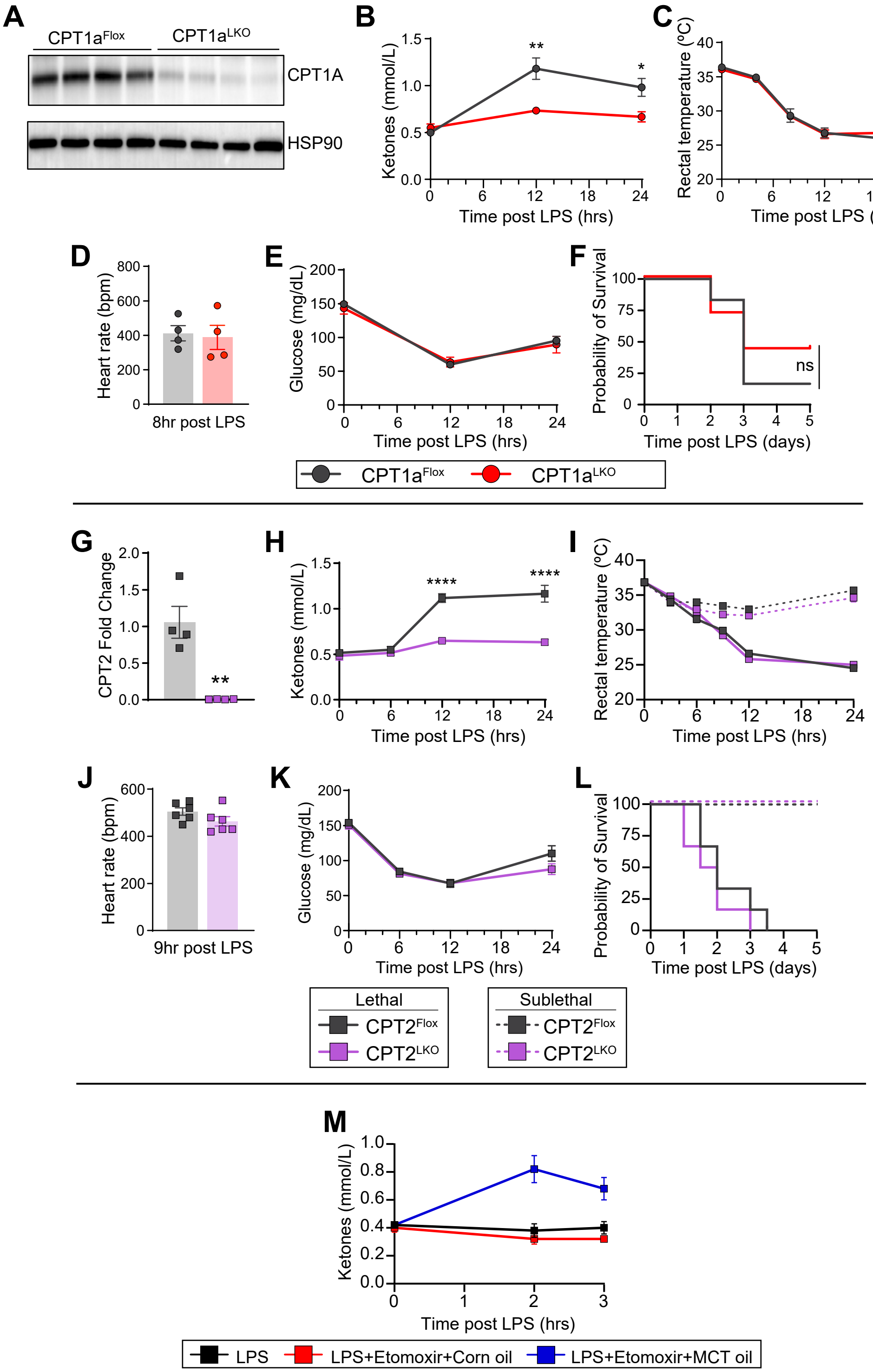

**Figure S6: Hepatic fatty acid oxidation is not required for tolerance to inflammation. (A)** Immunoblot of CPT1A in liver (n=4/group). Circulating **(B)** ketones (n=6-7/group), **(C)** rectal temperatures (n=6-7/group); and **(D)** heart rates (n=4/group), **(E)** glucose (n=6-7/group), and **(F)** Kaplan–Meier survival curve (n=6-7/group) after lipopolysaccharide (LPS) (5 mg/kg). **(G)** Expression of *Cpt2* in liver (n=4). Circulating **(H)** ketones, **(I)** rectal temperatures, **(J)** heart rates, **(K)** glucose, and **(L)** Kaplan–Meier survival curve after LPS. **(M)** Circulating ketones after LPS (5 mg/kg) and etomoxir with medium-chain triglyceride (MCT) or corn oil supplementation (n=5/group). For (**I** and **L**), LPS dose was 5 mg/kg (lethal) and 2.5 mg/kg (Sublethal) (n=6/group). For H, J, and K LPS dose was 5 mg/kg. Data are represented as mean ± SEM. *p < 0.05; **p < 0.01; ***p < 0.001; ****p < 0.0001, log-rank (Mantel–Cox) test.

**Figure S7: CL316,243 promotes tolerance through the adipose β3 receptor.**

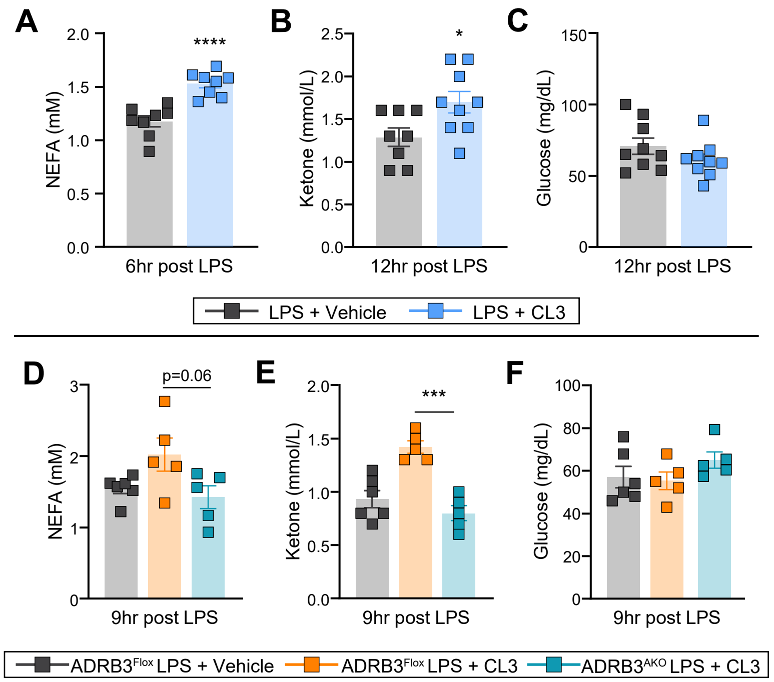

**Figure S7: CL316,243 promotes tolerance through the adipose β3 receptor.** Circulating **(A)** non-esterified fatty acids (NEFAs), **(B)** ketones, and **(C)** glucose levels after lipopolysaccharide (LPS) and CL316,243 (CL3). For (A-C) n=8/group. Circulating **(D)** NEFAs, **(E)** ketones, and **(F)** glucose levels after LPS and CL3. For (D-F), n=5-6/group. LPS dose was 10 mg/kg. Data are represented as mean ± SEM. *$p < 0.05$; **$p < 0.01$; ***$p < 0.001$; ****$p < 0.0001$.

**Figure S8: Glucose dissolved in water increases mortality with LPS.**

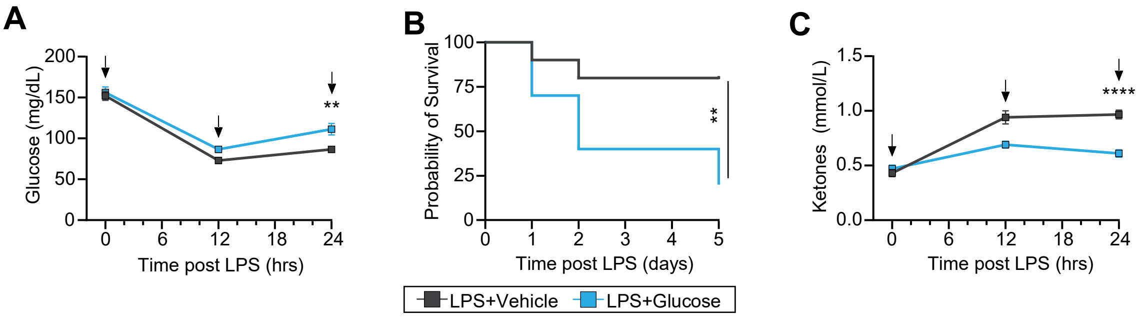

**Figure S8: Glucose dissolved in water increases mortality with LPS. (A)** Glucose, **(B)** Kaplan–Meier survival curve and **(C)** ketones after lipopolysaccharide (LPS) with glucose supplementation. Data are represented as mean ± SEM. *$p < 0.05$; **$p < 0.01$; ***$p < 0.001$; ****$p < 0.0001$, log-rank (Mantel–Cox) test. LPS dose was 10 mg/kg and n=10/group. Arrows indicate the time of CL3 administration and intralipid supplementation. Veh, vehicle. Vehicle was PBS, and glucose was dissolved in water.

**Supplemental Table 1. Encounter characteristics of control patients and those exposed to a β3-agonist before and after propensity score matching.**

| Encounter Characteristics | Exposed (n=14,345) | Unexposed Unmatched (n=3,953,862) | Unexposed PS-Matched (n=14,345) | SMD |
|---|---|---|---|---|
| Length of Stay (Days), mean (SD) | 7.1 (4.8) | 8.0 (5.5) | 7.2 (4.7) | — |
| Pressor Use | 657 (4.6%) | 471,426 (11.9%) | 650 (4.5%) | 0.0023 |
| Pressor-Free Days, mean (SD) | 8.8 (6.0) | 9.3 (6.5) | 8.7 (5.9) | — |
| % Stay Pressor-Free, mean (SD) | 83.8 (18.5) | 79.3 (23.3) | 82.0 (21.5) | — |
| ICU Length of Stay (Days), mean (SD) | 5.2 (6.6) | 5.0 (4.7) | 4.0 (3.6) | — |
| Ventilator Use During Admission | 443 (3.1%) | 475,774 (12.0%) | 420 (2.9%) | 0.0094 |
| **Average First 24-hour Vitals** | | | | |
| Temperature (F) , mean (SD) | 98.2 (0.9) | 98.2 (1.0) | 98.2 (1.0) | 0.0893 |
| Respiratory Rate (/min) , mean (SD) | 18.6 (2.8) | 19.0 (3.4) | 18.6 (2.8) | -0.0013 |
| Pulse Rate (/min) , mean (SD) | 83.1 (14.4) | 86.2 (16.0) | 82.9 (14.7) | 0.0129 |
| Systolic Blood Pressure (mmHg) , mean (SD) | 129.6 (19.1) | 126.9 (20.2) | 129.3 (19.7) | 0.0180 |
| Diastolic Blood Pressure (mmHg) , mean (SD) | 68.9 (10.8) | 70.0 (11.8) | 68.8 (11.0) | 0.0076 |
| Oxygen Saturation (%) , mean (SD) | 95.1 (2.3) | 95.6 (2.5) | 95.1 (2.7) | 0.0047 |

**Table S1. Encounter characteristics of control patients and those exposed to a β3-agonist before and after propensity score matching.** ICU, intensive care unit; SMD, standardized mean difference; PS, propensity score.
